# Endogenous hydrogen peroxide positively regulates secretion of a gut-derived peptide in neuroendocrine potentiation of the oxidative stress response in *C. elegans*

**DOI:** 10.1101/2024.04.03.587937

**Authors:** Qi Jia, Drew Young, Qixin Zhang, Derek Sieburth

## Abstract

The gut-brain axis mediates bidirectional signaling between the intestine and the nervous system and is critical for organism-wide homeostasis. Here we report the identification of a peptidergic endocrine circuit in which bidirectional signaling between neurons and the intestine potentiates the activation of the antioxidant response in *C. elegans* in the intestine. We identify a FMRF-amide-like peptide, FLP-2, whose release from the intestine is necessary and sufficient to activate the intestinal oxidative stress response by promoting the release of the antioxidant FLP-1 neuropeptide from neurons. FLP-2 secretion from the intestine is positively regulated by endogenous hydrogen peroxide (H_2_O_2_) produced in the mitochondrial matrix by *sod-3*/superoxide dismutase, and is negatively regulated by *prdx-2*/peroxiredoxin, which depletes H_2_O_2_ in both the mitochondria and cytosol. H_2_O_2_ promotes FLP-2 secretion through the DAG and calcium-dependent protein kinase C family member *pkc-2* and by the SNAP25 family member *aex-4* in the intestine. Together, our data demonstrate a role for intestinal H_2_O_2_ in promoting inter-tissue antioxidant signaling through regulated neuropeptide-like protein exocytosis in a gut-brain axis to activate the oxidative stress response.

## Introduction

The gut-brain axis is critical for communication between the intestine and the nervous system to regulate behavior and maintain homeostasis, and altered gut-brain signaling is associated with neurodegeneration, obesity and tumor proliferation (Carabotti et al. 2015; Grenham et al. 2011; Mayer, Nance, and Chen 2022; Mehrian-Shai et al. 2019; Vitali et al. 2022). Over the last decade the importance of peptides that function as signals in gut-brain signaling has gained recognition. Numerous gut peptides are distributed throughout the gastrointestinal (GI) tract with regional specificity (Haber et al. 2017), and gut-secreted peptides can modulate neurocircuits that regulate feeding behavior and glucose metabolism (Batterham and Bloom 2003; Han et al. 2018; Song et al. 2019), inflammatory responses against pathogenic bacteria (Campos-Salinas et al. 2014; Yu et al. 2021), and satiety (Batterham et al. 2002; Chelikani, Haver, and Reidelberger 2005; Gibbs, Young, and Smith 1973; Lutz et al. 1995; Lutz, Del Prete, and Scharrer 1994; West, Fey, and Woods 1984). A gut-released peptide suppresses arousal through dopaminergic neurons during sleep in *Drosophila* (Titos et al. 2023). In *C. elegans*, gut-derived peptides regulate rhythmic behavior and behavioral responses to pathogenic bacteria (Lee and Mylonakis 2017; Singh and Aballay 2019; Wang et al. 2013). Conversely, peptides released from the nervous system regulate many aspects of intestinal function including gut mobility, inflammation and immune defense (Browning and Travagli 2014; Furness et al. 2014; Lai, Mills, and Chiu 2017). In *C. elegans*, the secretion of peptides from various neurons regulates the mitochondrial unfolded protein response (UPR^mt^), the heat shock response, and the antioxidant response in the intestine (Jia and Sieburth 2021; Maman et al. 2013; Prahlad, Cornelius, and Morimoto 2008; Shao, Niu, and Liu 2016). In spite of the many roles of peptides in the gut-brain axis, the mechanisms underlying the regulation of intestinal peptide secretion and signaling remain to be fully defined.

Hydrogen peroxide (H_2_O_2_) is emerging as an important signaling molecule that regulates intracellular signaling pathways by modifying specific reactive residues on target proteins. For example, H_2_O_2_-regulated phosphorylation of inhibitor of nuclear factor κB (NF-κB) kinase, leads to the activation of NF-κB during development, inflammation and immune responses. (Kamata et al. 2002; Oliveira-Marques et al. 2009; Takada et al. 2003). In addition, H_2_O_2_-induced tyrosine and cysteine modifications contribute to redox regulation of c-Jun N-terminal kinase 2 (JNK2), Src family kinase, extracellular signal-regulated kinases 1 and 2 (ERK1/2), protein kinase C (PKC) and other protein kinases (Kemble and Sun 2009; Konishi et al. 1997; Lee et al. 2003; Nelson et al. 2018). H_2_O_2_ signaling has been implicated in regulating neurotransmission and transmitter secretion. H_2_O_2_ at low concentration increases neurotransmission at neuromuscular junctions without influencing lipid oxidation (A R Giniatullin and R A Giniatullin 2003; Giniatullin, Petrov, and Giniatullin 2019; Shakirzyanova et al. 2009), and enhanced endogenous H_2_O_2_ generation regulates dopamine release (Avshalumov et al. 2005; Avshalumov and Rice 2003; Bao, Avshalumov, and Rice 2005; Chen, Avshalumov, and Rice 2001, 2002). Acute H_2_O_2_ treatment increases exocytosis of ATP-containing vesicles in astrocytes (Li et al. 2019). Finally, mitochondrially-derived H_2_O_2_ regulates neuropeptide release from neurons in *C. elegans* (Jia and Sieburth 2021). Cellular H_2_O_2_ levels are tightly controlled through the regulation of its production from superoxide by superoxide dismutases (SODs) and cytoplasmic oxidases (Fridovich 1995, 1997; Messner and Imlay 2002; Zelko, Mariani, and Folz 2002), and through its degradation by catalases, peroxidases, and peroxiredoxins (Chance, Sies, and Boveris 1979; Marinho et al. 2014). In the intestine, endogenously produced H_2_O_2_ plays important roles as an antibacterial agent in the lumen, and in activating the ER unfolded protein response (UPR^ER^) through protein sulfenylation (Botteaux et al. 2009; Corcionivoschi et al. 2012; Hourihan et al. 2016; Miller et al. 2020).

Here we demonstrate a role of endogenous H_2_O_2_ signaling in the intestine in regulating the release of the intestinal FMRFamide-like peptide, FLP-2, to modulate a neurocircuit that activates the anti-oxidant response in the intestine in *C. elegans.* Intestinal FLP-2 signaling functions by potentiating the release of the antioxidant neuropeptide-like protein FLP-1 from AIY interneurons, which in turn activates the antioxidant response in the intestine. FLP-2 secretion from the intestine is rapidly and positively regulated by H_2_O_2_, whose levels are positively regulated by superoxide dismutases in the mitochondrial matrix and cytosol, and negatively regulated by the peroxiredoxin-thioredoxin system in the cytosol. Intestinal FLP-2 release is mediated by *aex-4*/SNAP25-dependent exocytosis of dense core vesicles and H_2_O_2_-induced FLP-2 secretion is dependent upon the production of intestinal diacylglycerol and on *pkc-2*/PKCα/β kinase activity.

## Results

### Neuronal FLP-1 secretion is regulated by neuropeptide signaling from the intestine

We previously showed that 10 minute treatment with the mitochondrial toxin juglone leads to a rapid, reversible and specific increase in FLP-1 secretion from AIY, as measured by a two-fold increase in coelomocyte fluorescence in animals expressing FLP-1::Venus fusion proteins in AIY (Fig. 1A and B, (Jia and Sieburth 2021)). Coelomocytes take up secreted neuropeptides by bulk endocytosis (Fares and Greenwald 2001) and the fluorescence intensity of Venus in their endocytic vacuoles is used as a measure of regulated neuropeptide secretion efficacy (Ailion et al. 2014; Ch’ng, Sieburth, and Kaplan 2008; Sieburth, Madison, and Kaplan 2006). To determine the role of the intestine in regulating FLP-1 secretion, we first examined *aex-5* mutants. *aex-5* encodes an intestinal subtilisin/kexin type 5 prohormone convertase that functions to proteolytically process peptide precursors into mature peptides in dense core vesicles (DCVs) (Edwards et al. 2019; Thacker and Rose 2000), and *aex-5* mutants are defective in peptide signaling from the intestine (Mahoney et al. 2008). We found that *aex-5* mutants expressing FLP-1::Venus in AIY exhibited no significant difference in coelomocyte fluorescence compared to wild type controls in the absence of juglone (Fig. 1B). However, coelomocyte fluorescence did not significantly increase in *aex-5* mutants treated with juglone. Expression of *aex-5* cDNA selectively in the intestine (under the *ges-1* promoter) fully restored normal responses to juglone to *aex-5* mutants, whereas *aex-5* cDNA expression in the nervous system (under the *rab-3* promoter) failed to rescue (Fig. 1B). Thus, peptide processing in intestinal DCVs is necessary for juglone-induced FLP-1 secretion from AIY.

**Figure 1.**
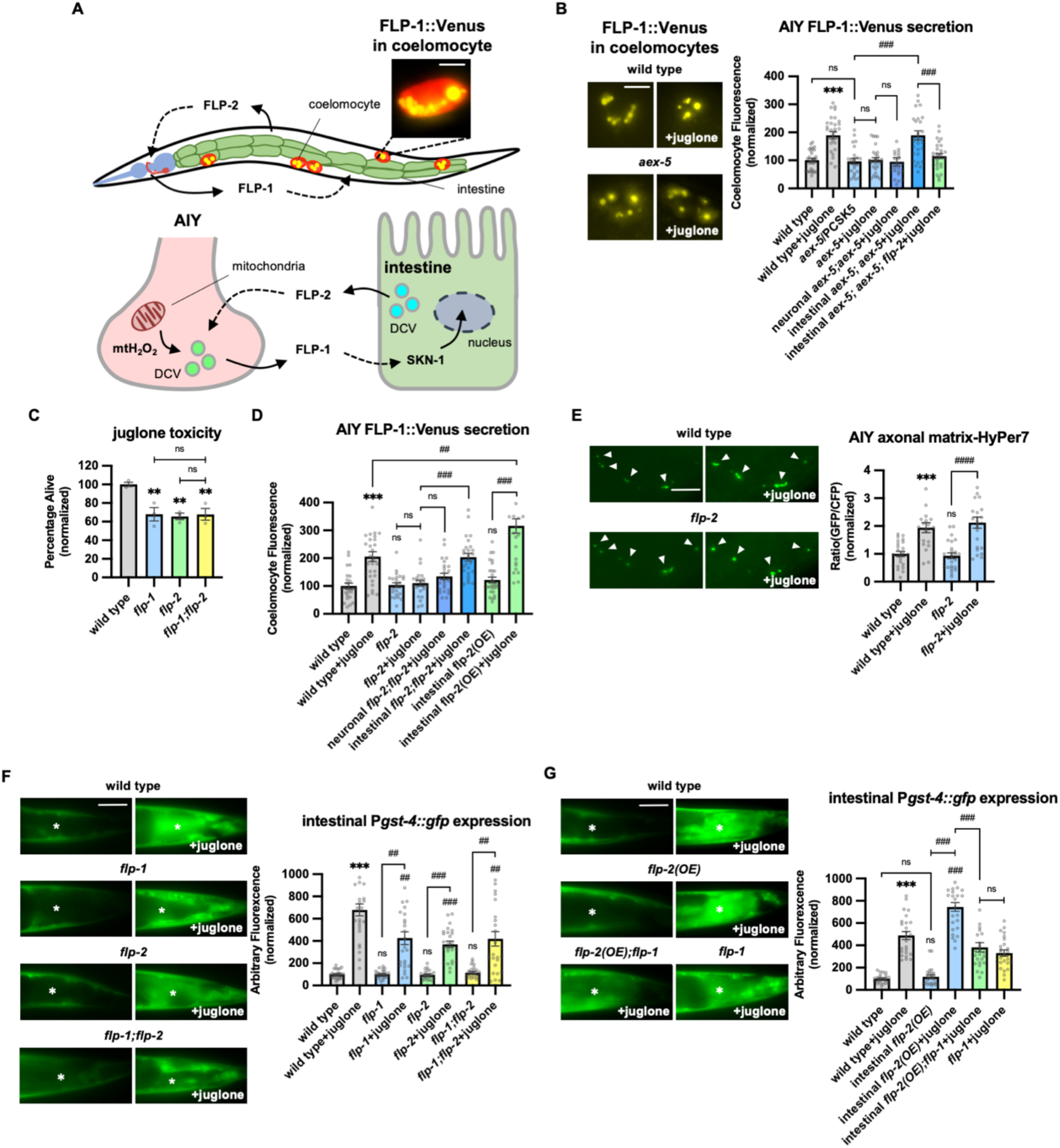
Peptidergic gut-to-neuron FLP-2 signaling potentiates the oxidative stress response. **A** (Top) Schematic showing the positions of AIY, intestine and coelomocytes of transgenic animals co-expressing FLP-1::Venus in the intestine and mCherry in coelomocytes. Representative image of the posterior coelomocyte that has taken up Venus into the endocytic compartment. Scale bar: 5μM. (Bottom) Schematic showing FLP-1 and FLP-2 peptides as inter-tissue signals in gut-intestine regulation of the antioxidant response. **B** Representative images and quantification of average coelomocyte fluorescence of the indicated mutants expressing FLP-1::Venus fusion proteins in AIY following M9 or 300μM juglone treatment for 10min. Neuronal *aex-5* denotes expression of *aex-5* cDNA under the *rab-3* promoter; intestinal *aex-5* denotes expression of *aex-5* cDNA under the *ges-1* promoter. Unlined *** denotes statistical significance compared to “wild type”. n = 30, 30, 24, 30, 26, 30, 30 independent animals. Scale bar: 5μM. **C** Average percentage of surviving young adult animals of the indicated genotypes after 16h recovery following 4h juglone treatment. Unlined ** denotes statistical significance compared to “wild type”. n = 213, 156, 189, 195 independent biological samples over three independent experiments. **D** Quantification of average coelomocyte fluorescence of the indicated mutants expressing FLP-1::Venus fusion proteins in AIY following M9 or 300μM juglone treatment for 10min. Neuronal *flp-2* denotes expression of *flp-2* gDNA under the *rab-3* promoter; intestinal *flp-2* denotes expression of *flp-2* gDNA under the *ges-1* promoter; intestinal *flp-2(OE)* denotes expression of *flp-2* gDNA under the *ges-1* promoter in wild type animals. Unlined *** and ns denote statistical significance compared to “wild type”. n = 20, 20, 25, 20, 20, 20, 25, 22 independent animals. **E** Representative images and quantification of fluorescence of mitochondrial matrix-targeted HyPer7 in the axon of AIY following M9 or 300μM juglone treatment for 10min. Arrowheads denote puncta marked by mito::HyPer7 fusion proteins (Excitation: 500 and 400nm; emission: 520nm). Ratio of images taken with 500nM (GFP) and 400nM (CFP) for excitation was used to measure H_2_O_2_ levels. Unlined *** and ns denote statistical significance compared to “wild type”. n = 24, 22, 25, 24 independent animals. Scale bar: 10μM. **F** Representative images and quantification of average fluorescence in the posterior intestine of transgenic animals expressing P*gst-4::gfp* after 1h M9 or juglone exposure and 3h recovery. Asterisks mark the intestinal region used for quantification. P*gst-4::gfp* expression in the body wall muscles, which appears as fluorescence on the edge animals in some images, was not quantified. Unlined *** and ns denote statistical significance compared to “wild type”; unlined ## and ### denote statistical significance compared to “wild type+juglone”. n = 25, 26, 25, 25, 25, 25, 25, 25 independent animals. Scale bar: 10μM. **G** Representative images and quantification of average fluorescence in the posterior region of transgenic animals expressing P*gst-4::gfp* after 1h M9 or juglone exposure and 3h recovery. Asterisks mark the intestinal region for quantification. P*gst-4::gfp* expression in the body wall muscles, which appears as fluorescence on the edge animals in some images, was not quantified. Unlined *** denotes statistical significance compared to “wild type”; unlined ### denotes statistical significance compared to “wild type+juglone”. n = 23, 25, 25, 26, 24, 25 independent animals. Scale bar: 10μM. **B-G** Data are mean values ± s.e.m normalized to wild type controls. ns. not significant, ** and ## *P <* 0.01, *** and ### *P* < 0.001 by Brown-Forsythe and Welch ANOVA with Dunnett’s T3 multiple comparisons test.

Next, we examined a number of mutants with impaired SNARE-mediated vesicle release in the intestine including *aex-1*/UNC13, *aex-*3/MADD, aex*-4*/SNAP25b, and *aex-6*/Rab27 (Fig. 2C (Iwasaki et al., 1997; Mahoney et al., 2006; Thacker & Rose, 2000; Thomas, 1990; Wang et al., 2013)), and they each exhibited no increases in FLP-1 secretion following juglone treatment above levels observed in untreated controls (Fig. S1A). NLP-40 is a neuropeptide-like protein whose release from the intestine is presumed to be controlled by *aex-1*, *aex-3*, *aex-4* and *aex-6* (Lin-Moore, Oyeyemi, and Hammarlund 2021; Mahoney et al. 2008; Shi et al. 2022; Wang et al. 2013). Null mutants in *nlp-40* or its receptor, *aex-2* (Wang et al. 2013), exhibited normal juglone-induced FLP-1 secretion (Fig. S1B). These results establish a gut-to-neuron signaling pathway that regulates FLP-1 secretion from AIY that is likely to be controlled by peptidergic signaling distinct from NLP-40.

**Figure 2.**
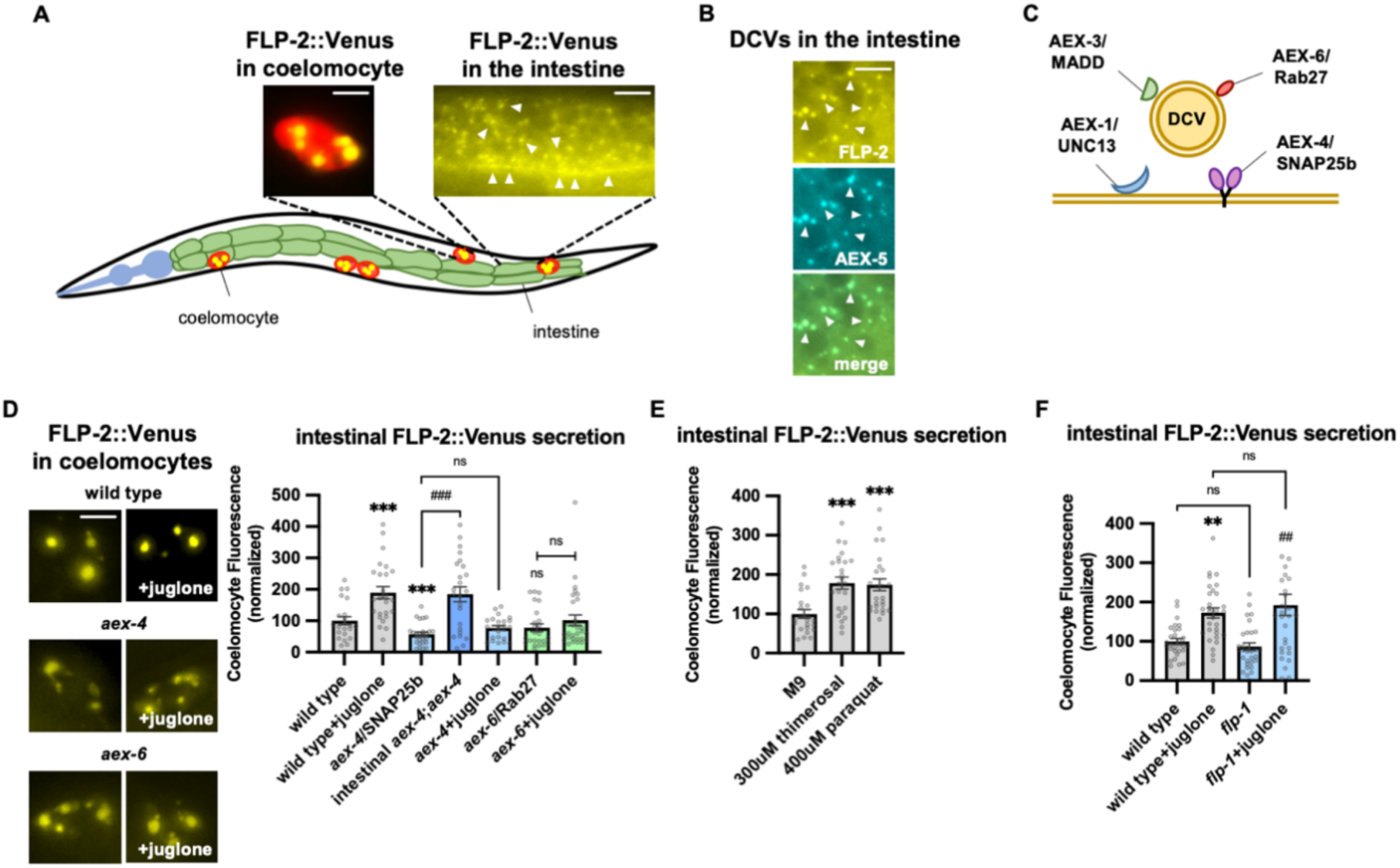
FLP-2 secretion from the intestine is stress regulated. **A** Schematic showing the positions of intestine and coelomocytes of transgenic animals co-expressing FLP-2::Venus in the intestine and mCherry in coelomocytes. Representative images of the posterior coelomocyte that have taken up Venus into the endocytic compartment (Scale bar: 5μM) and the posterior intestinal region showing the distribution of FLP-2::Venus in puncta in the intestine are shown (Scale bar: 15μM). **B** Representative images of fluorescence distribution in the posterior intestinal region of transgenic animals co-expressing FLP-2::Venus and AEX-5::mTur2 fusion proteins. Arrowheads denote puncta marked by both fusion proteins. Scale bar: 5μM. **C** Schematic showing the locations of AEX-1/UNC13, AEX-3/MADD, AEX-4/SNAP25 and AEX-6/Rab27 relative to a DCV. **D** Representative images and quantification of average coelomocyte fluorescence of the indicated mutants expressing FLP-2::Venus fusion proteins in the intestine following M9 or 300μM juglone for 10min. Unlined *** and ns denote statistical significance compared to “wild type”. n = 29, 25, 24, 30, 23, 30, 25, 25, 25 independent animals. Scale bar: 5μM. **E** Quantification of average coelomocyte fluorescence of transgenic animals expressing FLP-2::Venus fusion proteins in the intestine following treatment with M9 buffer or the indicated stressors for 10min. Unlined *** denotes statistical significane compared to “M9”. n = 23, 25, 25 independent animals. **F** Quantification of average coelomocyte fluorescence of the indicated mutants expressing FLP-2::Venus fusion proteins in the intestine following M9 or 300μM juglone treatment for 10min. Unlined ** denotes statistical significance compared to “wild type”; unlined ## denotes statistical significance compared to “*flp-1”*; a denotes statistical significance compared to “wild type+juglone”. n = 30, 30, 30, 30 independent animals. **D-F** Data are mean values ± s.e.m normalized to wild type controls. ns. not significant, ** and ## *P <* 0.01, *** and *### P <* 0.001 by Brown-Forsythe and Welch ANOVA with Dunnett’s T3 multiple comparisons test.

### FLP-2 signaling from the intestine potentiates neuronal FLP-1 secretion and the oxidative stress response

*flp-1* protects animals from the toxic effects of juglone (Jia and Sieburth 2021). We reasoned that the intestinal signal that regulates FLP-1 secretion should also protect animals from juglone-induced toxicity. We identified the FMRF-amide neuropeptide-like protein, *flp-2*, in an RNA interference (RNAi) screen for neuropeptides that confer hypersensitivity to juglone toxicity upon knockdown (Jia and Sieburth 2021). *flp-2* signaling has been implicated in regulating lifespan, reproductive development, locomotion during lethargus, and the mitochondrial unfolded protein response (UPR^mt^) (Chai et al. 2022; Chen et al. 2016; Kageyama et al. 2022; Shao et al. 2016). Putative *flp-2(ok3351)* null mutants, which eliminate most of the *flp-2* coding region, are superficially as healthy as wild type animals, but they exhibited significantly reduced survival in the presence of juglone compared to wild type controls (Fig. 1C). The reduced survival rate of *flp-2* mutants was similar to that of *flp-1* mutants, and *flp-1; flp-2* double mutants exhibited survival rates that were not more severe than those of single mutants (Fig. 1C), suggesting that *flp-1* and *flp-2* may function in a common genetic pathway.

To determine whether *flp-2* signaling regulates FLP-1 secretion from AIY, we examined FLP-1::Venus secretion. *flp-2* mutants exhibited normal levels of FLP-1 secretion in the absence of stress, but FLP-1 secretion failed to significantly increase following juglone treatment of *flp-2* mutants (Fig. 1D). *flp-2* is expressed in a subset of neurons as well as the intestine (Chai et al. 2022), and *flp-2* functions from the nervous system for its roles in development and the UPR^mt^ (Chai et al. 2022; Chen et al. 2016; Kageyama et al. 2022; Shao et al. 2016). Expressing a *flp-2* genomic DNA, fragment (containing both the *flp-2a* and *flp-2b* isoforms that arise by alternative splicing), specifically in the nervous system failed to rescue the FLP-1::Venus defects of *flp-2* mutants, whereas expressing *flp-2* selectively in the intestine fully restored juglone-induced FLP-1::Venus secretion to *flp-2* mutants (Fig. 1D). These results indicate that *flp-2* signaling is dispensable for FLP-1 secretion from AIY under normal conditions, but that *flp-2* originating from the intestine is necessary to increase FLP-1 secretion during oxidative stress.

To address how *flp-2* signaling regulates FLP-1 secretion from AIY, we examined H_2_O_2_ levels in AIY using a mitochondrially targeted pH-stable H_2_O_2_ sensor HyPer7 (mito-HyPer7, Pak et al. 2020). Mito-HyPer7 adopted a punctate pattern of fluorescence in AIY axons, and the average fluorescence intensity of axonal mito-HyPer7 puncta increased about two-fold following 10 minute juglone treatment (Fig 1E), in agreement with our previous studies using HyPer (Jia and Sieburth 2021), confirming that juglone rapidly increases mitochondrial AIY H_2_O_2_ levels. *flp-2* mutations had no significant effects on the localization or the average intensity of mito-HyPer7 puncta in AIY axons either in the absence of juglone, or in the presence of juglone (Fig 1E), suggesting that *flp-2* signaling promotes FLP-1 secretion by a mechanism that does not increase H_2_O_2_ levels in AIY. Consistent with this, intestinal overexpression of *flp-*2 had no effect on FLP-1::Venus secretion in the absence of juglone, but significantly enhanced the ability of juglone to increase FLP-1 secretion (Fig. 1D). We conclude that both elevated mitochondrial H_2_O_2_ levels and intact *flp-2* signaling from the intestine are necessary to increase FLP-1 secretion from AIY.

Previously we showed that FLP-1 signaling from AIY positively regulates the activation of the antioxidant transcription factor SKN-1/Nrf2 in the intestine. Specifically, *flp-1* mutations impair the juglone-induced expression of the SKN-1 reporter transgene P*gst-4::gfp* (Fig. 1F and (Jia and Sieburth 2021)). We found that mutations in *flp-2* caused a similar reduction in juglone-induced P*gst-4::gfp* expression as *flp-1* mutants, and that *flp-1; flp-2* double mutants exhibited similar impairments in juglone-induced P*gst-4::gfp* expression as *flp-1* or *flp-2* single mutants (Fig. 1F). Conversely, overexpression of *flp-2* selectively in the intestine elevated juglone-induced P*gst-4::gfp* expression, without altering baseline P*gst-4::gfp* expression, and the elevated P*gst-4::gfp* expression in juglone-treated animals overexpressing *flp-2* was entirely dependent upon *flp-1* (Fig. 1G). It’s noteworthy that overexpressing *flp-2* in the intestine did not enhance FLP-1::Venus release or *Pgst-4::gfp* expression in the absence of stress, indicating that the FLP-1 mediated anti-oxidant pathway by *flp-2* is stress-activated. Together this data indicates that *flp-2* signaling originating in the intestine positively regulates the stress-induced secretion of FLP-1 from AIY, as well as the subsequent activation of anti-oxidant response genes in the intestine. We propose that FLP-1 and FLP-2 define a bidirectional gut-neuron signaling axis, whereby during periods of oxidative stress, FLP-2 released from the intestine positively regulates FLP-1 secretion from AIY, and FLP-1, in turn, potentiates the antioxidant response in the intestine (Fig. 1A).

### FLP-2 secretion from the intestine is H_2_O_2_-regulated

To directly investigate the mechanisms underlying the regulation of FLP-2 secretion, we examined FLP-2::Venus fusion proteins expressed in the intestine under various conditions (Fig. 2A). FLP-2::Venus fusion proteins adopted a punctate pattern of fluorescence throughout the cytoplasm of intestinal cells and at the plasma membrane (Fig. 2A), and FLP-2::Venus puncta co-localized with the DCV cargo protein AEX-5/PCSK5 tagged to mTurquoise2 (AEX-5::mTur2, Fig. 2B). FLP-2::Venus fluorescence was also observed in the coelomocytes (marked by mCherry) (Fig. 2A, D), indicating that FLP-2 is released from the intestine. SNAP25 forms a component of the core SNARE complex, which drives vesicular membrane fusion and transmitter release (Chen and Scheller 2001; Goda 1997; Jahn and Scheller 2006). *aex-4* encodes the *C. elegans* homolog of SNAP25, and mutations in *aex-4* disrupt the secretion of neuropeptides from the intestine (Lin-Moore et al. 2021; Mahoney et al. 2008; Wang et al. 2013) (Fig. 2C). We found that *aex-4* null mutations significantly reduced coelomocyte fluorescence in FLP-2::Venus expressing animals, and expression of *aex-4* cDNA selectively in the intestine fully restored FLP-2 secretion to *aex-4* mutants (Fig. 2D). Together these results suggest that intestinal FLP-2 can be packaged into DCVs that undergo release via SNARE-dependent exocytosis.

To test whether intestinal FLP-2 secretion is regulated by oxidative stress, we examined coelomocyte fluorescence in FLP-2::Venus-expressing animals that had been exposed to a number of different commonly used oxidative stressors. We found that 10 minute exposure to juglone, thimerosal, or paraquat, which promote mitochondria-targeted toxicity (Castello, Drechsel, and Patel 2007; Elferink 1999; Sharpe, Livingston, and Baskin 2012), each significantly increased Venus fluorescence intensity in the coelomocytes compared to untreated controls (Fig. 2D and E). We conducted four controls for specificity: First, juglone treatment did not significantly alter fluorescence intensity of mCherry expressed in coelomocytes (Fig. S2A). Second, impairing intestinal DCV secretion (by either *aex-4*/SNAP25 or *aex-6*/Rab27 mutations (Lin-Moore et al. 2021; Mahoney et al. 2006, 2008; Thomas 1990), blocked the juglone-induced increase in coelomocyte fluorescence in FLP-2::Venus-expressing animals (Fig. 2C). Third, *nlp-40* and *nlp-27* encode neuropeptide-like proteins that are released from the intestine but are not implicated in stress responses (Liu et al. 2023; Taylor et al. 2021; Wang et al. 2013). Juglone treatment had no detectable effects on coelomocyte fluorescence in animals expressing intestinal NLP-40::Venus or NLP-27::Venus fusion proteins (Fig. S2B and C), and NLP-40::mTur2 puncta did not overlap with FLP-2::Venus puncta in the intestine (Fig. S2D). Finally, *flp-1* mutants exhibited wild type levels of FLP-2 secretion both in the absence and presence of juglone (Fig. 2E). The distribution of FLP-2::Venus puncta in the intestine was not detectably altered by juglone treatment. Together, these results indicate that acute oxidative stress selectively increases the exocytosis of FLP-2-containing DCVs from the intestine, upstream of *flp-1* signaling.

### SOD-1 and SOD-3 superoxide dismutases regulate FLP-2 release

Juglone generates superoxide anion radicals (Ahmad and Suzuki 2019; Paulsen and Ljungman 2005) and juglone treatment of *C. elegans* increases ROS levels (de Castro, Hegi de Castro, and Johnson 2004) likely by promoting the global production of mitochondrial superoxide. Superoxide can then be rapidly converted into H_2_O_2_ by superoxide dismutase. To determine whether H_2_O_2_ impacts FLP-2 secretion, we first examined superoxide dismutase mutants. *C. elegans* encodes five superoxide dismutase genes (*sod-1* through *sod-5*). *sod-1* or *sod-3* null mutations blocked juglone-induced FLP-2 secretion without altering baseline FLP-2 secretion, whereas *sod-2*, *sod-4*, or *sod-5* mutations had no effect on FLP-2 secretion in the presence of juglone (Fig. 3A and B, S3A). *sod-1; sod-3* double mutants exhibited juglone-induced FLP-2 secretion defects that was similar to single mutants, without significantly altering FLP-2 secretion in the absence of stress (Fig. 3C). *sod-1* encodes the ortholog of mammalian SOD1, which is a cytoplasmic SOD implicated in the development of amyotrophic lateral sclerosis (ALS) and cancer (Giglio et al. 1994; Papa, Manfredi, and Germain 2014; Wang et al. 2021; Zhang et al. 2007). SOD-1::fusion proteins adopted a diffuse pattern of fluorescence in intestinal cells, consistent with a cytoplasmic localization (Fig. 3D). Transgenes expressing the *sod-1* cDNA selectively in the intestine fully rescued the juglone-induced FLP-2::Venus secretion defects of *sod-1* mutants (Fig. 3A). *sod-3* encodes a homolog of mammalian SOD2, which is a mitochondrial matrix SOD implicated in protection against oxidative stress-induced neuronal cell death (Fukui and Zhu 2010; Vincent et al. 2007). Intestinal SOD-3::GFP fusion proteins localized to round structures that were surrounded by the outer membrane mitochondrial marker TOMM-20::mCherry (Ahier et al. 2018), consistent with a mitochondrial matrix localization (Fig. 3E). Expression of *sod-3* cDNA in the intestine fully restored juglone-induced FLP-2 release to *sod-3* mutants (Fig. 3B). *sod-3* variants lacking the mitochondrial localization sequence (*sod-3*(ΔMLS)), were no longer localized to mitochondria (Fig. 3F) and failed to restore normal responsiveness to juglone to *sod-3* mutants (Fig. 3B). Thus, the generation of H_2_O_2_ by either SOD-1 in the cytoplasm or by SOD-3 in the mitochondrial matrix is necessary for juglone to increase FLP-2 secretion.

**Figure 3.**
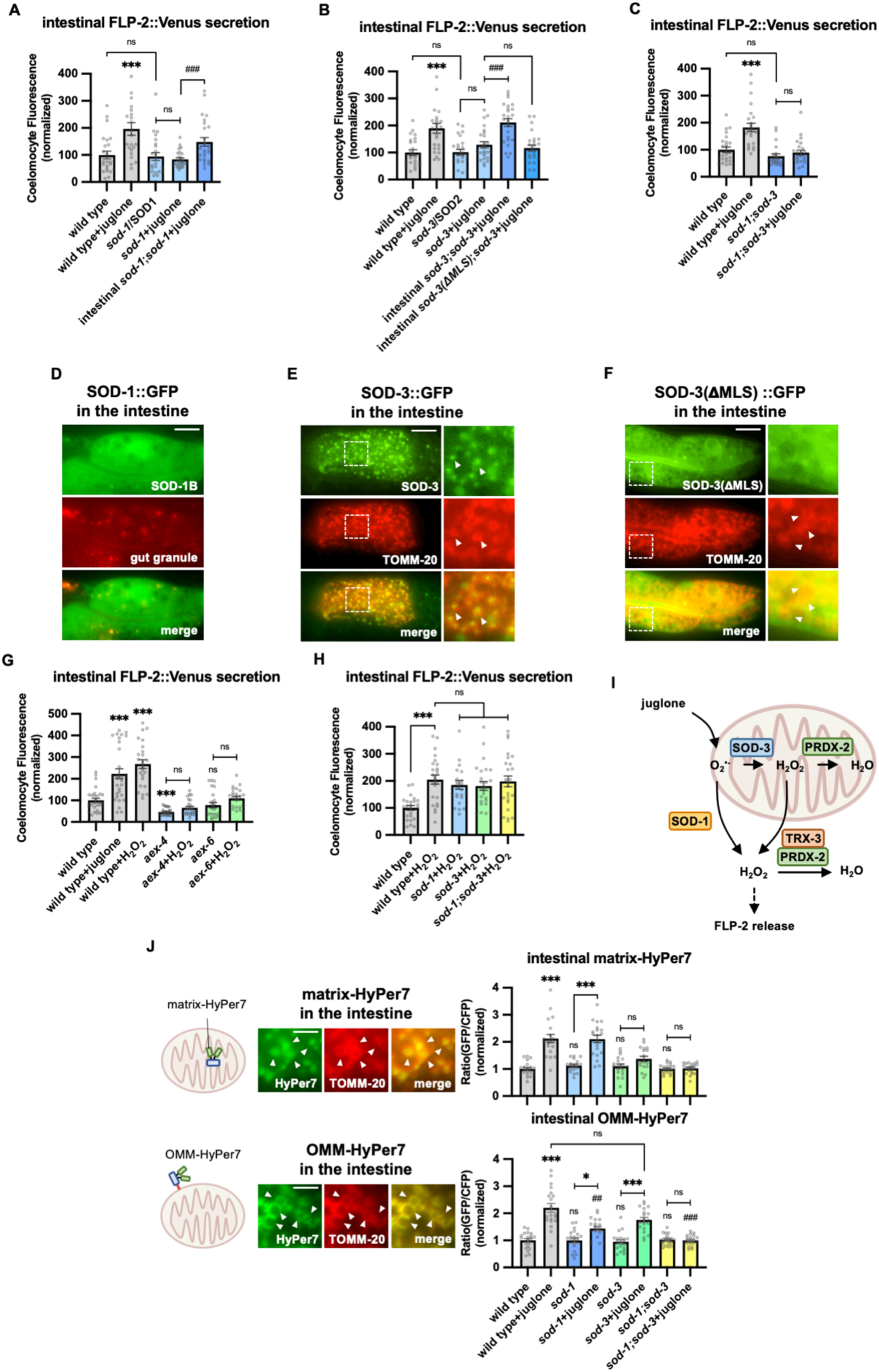
SOD-1/SOD-3 mediates endogenous H_2_O_2_ regulates FLP-2 release from the intestine. **A** Quantification of average coelomocyte fluorescence of the indicated mutants expressing FLP-2::Venus fusion proteins in the intestine following M9 or 300μM juglone treatment for 10min. Intestinal *sod-*1 denotes expression of *sod-1b* cDNA under the *ges-1* promoter. Unlined *** denotes statistical significance compared to “wild type”. n = 25, 22, 24, 24, 25 independent animals. **B** Quantification of average coelomocyte fluorescence of the indicated mutants expressing FLP-2::Venus fusion proteins in the intestine following M9 or 300μM juglone treatment for 10min. Intestinal *sod-3* and *sod-3(*ΔMLS) denote intestinal expression of *sod-3* cDNA and *sod-3(*ΔMLS) variants, which lacks the mitochondrial localization sequence, under the *ges-1* promoter. Unlined *** denotes statistical significance compared to “wild type”. n = 25, 25, 25, 25, 25, 25 independent animals. **C** Quantification of average coelomocyte fluorescence of the indicated mutants expressing FLP-2::Venus fusion proteins in the intestine following M9 or 300μM juglone treatment for 10min. Unlined *** denotes statistical significance compared to “wild type”. n = 25, 25, 22, 25 independent animals. **D** Representative images of fluorescence distribution in the posterior intestinal region of transgenic animals expressing SOD-1b::GFP fusion proteins in contrast against auto-fluorescence of gut granules. Scale bar: 10μM. **E** Representative images of fluorescence distribution in the posterior intestinal region of transgenic animals co-expressing SOD-3::GFP and TOMM-20::mCherry (to target mitochondria) fusion proteins. Scale bar: 15μM. **F** Representative images of fluorescence distribution in the posterior intestinal region of transgenic animals co-expressing SOD-3(ΔMLS)::GFP and TOMM-20::mCherry fusion proteins. Scale bar: 15μM. **G** Quantification of average coelomocyte fluorescence of the indicated mutants expressing FLP-2::Venus fusion proteins in the intestine following M9, 300μM juglone or 1mM H_2_O_2_ treatment for 10min. Unlined *** and ns denote statistical significance compared to “wild type”. n = 29, 30, 25, 25, 25, 24, 25 independent animals. **H** Quantification of average coelomocyte fluorescence of the indicated mutants expressing FLP-2::Venus fusion proteins in the intestine following M9 or 1mM H_2_O_2_ treatment for 10min. n = independent animals. **I** Schematic showing that SOD-1 and SOD-3 mediate juglone-induced H_2_O_2_ production in promoting FLP-2 release, and the PRDX-2/TRX-3 system detoxifies excessive H_2_O_2_. **J** Schematic, representative images and quantification of fluorescence in the posterior region of the indicated transgenic animals co-expressing mitochondrial matrix targeted HyPer7 (matrix-HyPer7) or mitochondrial outer membrane targeted HyPer7 (OMM-HyPer7) with TOMM-20::mCherry following M9 or 300μM juglone treatment. Ratio of images taken with 500nM (GFP) and 400nM (CFP) for excitation and 520nm for emission was used to measure H_2_O_2_ levels. Unlined *** and ns denote statistical significance compared to “wild type”. Unlined ## and ### denote statistical significance compared to “wild type+juglone”. (top) n = 20, 20, 18, 20, 19, 19, 20, 20 independent animals. (bottom) n = 20, 20, 19, 20, 20, 20, 20, 20 independent animals. Scale bar: 5μM. **A-C, G-H and J** Data are mean values ± s.e.m normalized to wild type controls. ns. not significant, * *P <* 0.05, ## *P <* 0.01, *** and ### *P <* 0.001 by Brown-Forsythe and Welch ANOVA with Dunnett’s T3 multiple comparisons test.

Next, to determine if H_2_O_2_ can regulate FLP-2 secretion, we treated animals acutely with exogenous H_2_O_2_. We found that 10 minute H_2_O_2_ treatment increased FLP-2::Venus secretion to a similar extent as 10 minute juglone treatment. *aex-4*/SNAP25, or *aex-6/*Rab27 mutants exhibited no increase in FLP-2 secretion in response to H_2_O_2_ treatment compared to untreated controls (Fig. 3G). In contrast, *sod-1* or *sod-3* mutants (or *sod-1; sod-3* double mutants) exhibited an increase in FLP-2 secretion in response to H_2_O_2_ that was similar to that of wild type controls (Fig. 3H), suggesting that exogenous H_2_O_2_ can bypass the requirement of SODs but not SNAREs to promote FLP-2 secretion. Together these results suggest that H_2_O_2_ generated by SODs can positively regulate intestinal FLP-2 exocytosis from DCVs (Fig. 3I).

### SOD-1 and SOD-3 regulate intestinal mitochondrial H_2_O_2_ levels

To directly monitor H_2_O_2_ levels in the intestine, we generated transgenic animals in which HyPer7 was targeted to either the mitochondrial matrix (matrix-HyPer7) by generating fusion proteins with the cytochrome c MLS, or to the cytosolic face of the outer mitochondrial membrane (OMM-HyPer7) by generating fusion proteins with TOMM-20. When co-expressed in the intestine with the OMM marker TOMM-20::mCherry, matrix-HyPer7 formed round structures throughout the cytoplasm that were surrounded by the OMM, and OMM-HyPer7 formed ring-like structures throughout the cytoplasm that co-localized with the OMM marker (Fig. 3J). Ten minute treatment with H_2_O_2_ significantly increased the fluorescence intensity by about two-fold of both matrix-HyPer7 and OMM-HyPer7 without altering mitochondrial morphology or abundance (Fig. 3J, S3B and C), validating the utility of HyPer7 as a sensor for acute changes in H_2_O_2_ levels in and around intestinal mitochondria.

Juglone treatment for 10 minutes led to a similar two-fold increase in matrix-HyPer7 fluorescence as H_2_O_2_ treatment (Fig. 3J). *sod-3* mutations did not alter baseline H_2_O_2_ levels in the matrix, but they completely blocked juglone-induced increases H_2_O_2_ levels, whereas *sod-1* mutations had no effect on either baseline or juglone-induced increases H_2_O_2_ levels (Fig. 3J). These results indicate that superoxide produced by juglone treatment is likely to be converted into H_2_O_2_ by SOD-3 in the mitochondrial matrix (Fig. 3I).

Next, we examined H_2_O_2_ levels on the outer surface of mitochondria using OMM-HyPer7 and we found that juglone treatment led to a two-fold increase in OMM-HyPer7 fluorescence, similar to H_2_O_2_ treatment (Fig. 3J). *sod-3* or *sod-1* mutations did not alter baseline H_2_O_2_ levels on the OMM, but *sod-1* single mutations attenuated, juglone-induced increases in OMM H_2_O_2_ levels, while *sod-3* mutations had no effect (Fig. 3J). In *sod-1; sod-3* double mutants, the juglone-induced increase in OMM H_2_O_2_ levels was completely blocked, whereas baseline H_2_O_2_ levels in the absence of stress were unchanged (Fig. 3J). These results suggest that *sod-3* and *sod-1* are exclusively required for H_2_O_2_ production by juglone and that both mitochondrial SOD-3 and cytosolic SOD-1 contribute to H_2_O_2_ levels in the cytosol. One model that could explain these results is that juglone-generated superoxide is converted into H_2_O_2_ both by SOD-3 in the matrix, and by SOD-1 in the cytosol, and that the H_2_O_2_ generated in the matrix can exit the mitochondria to contribute to cytosolic H_2_O_2_ levels needed to drive FLP-2 secretion (Fig. 3I).

### The peroxiredoxin-thioredoxin system regulates endogenous H_2_O_2_ levels and FLP-2 secretion

To determine whether endogenous H_2_O_2_ regulates FLP-2 secretion, we examined mutations in the peroxiredoxin-thioredoxin system. Peroxiredoxins and thioredoxins detoxify excessive H_2_O_2_ by converting it into water and they play a critical role in maintaining cellular redox homeostasis (Netto and Antunes 2016) (Fig. 4A). *C. elegans* encodes two peroxiredoxin family members, *prdx-2* and *prdx-3*, that are expressed at high levels in the intestine (Taylor et al. 2021). Null mutations in *prdx-2* significantly increased FLP-2::Venus secretion compared to wild type animals in the absence of stress (Fig. 4B), whereas null mutations in *prdx-3* had no effect on FLP-2 secretion (Fig. S4A). We observed a corresponding increase in both matrix-HyPer7 and OMM-HyPer7 fluorescence intensity in *prdx-2* mutants (Fig. 4C and D), demonstrating that endogenous H_2_O_2_ is neutralized by peroxiredoxin and establishing a correlation between endogenous H_2_O_2_ levels and FLP-2 secretion. The increase in FLP-2 secretion in *prdx-2* mutants was not further increased by juglone treatment (Fig. 4B). These results suggest that elevation in the levels of endogenously produced H_2_O_2_ in the intestine can positively regulate FLP-2 secretion.

**Figure 4.**
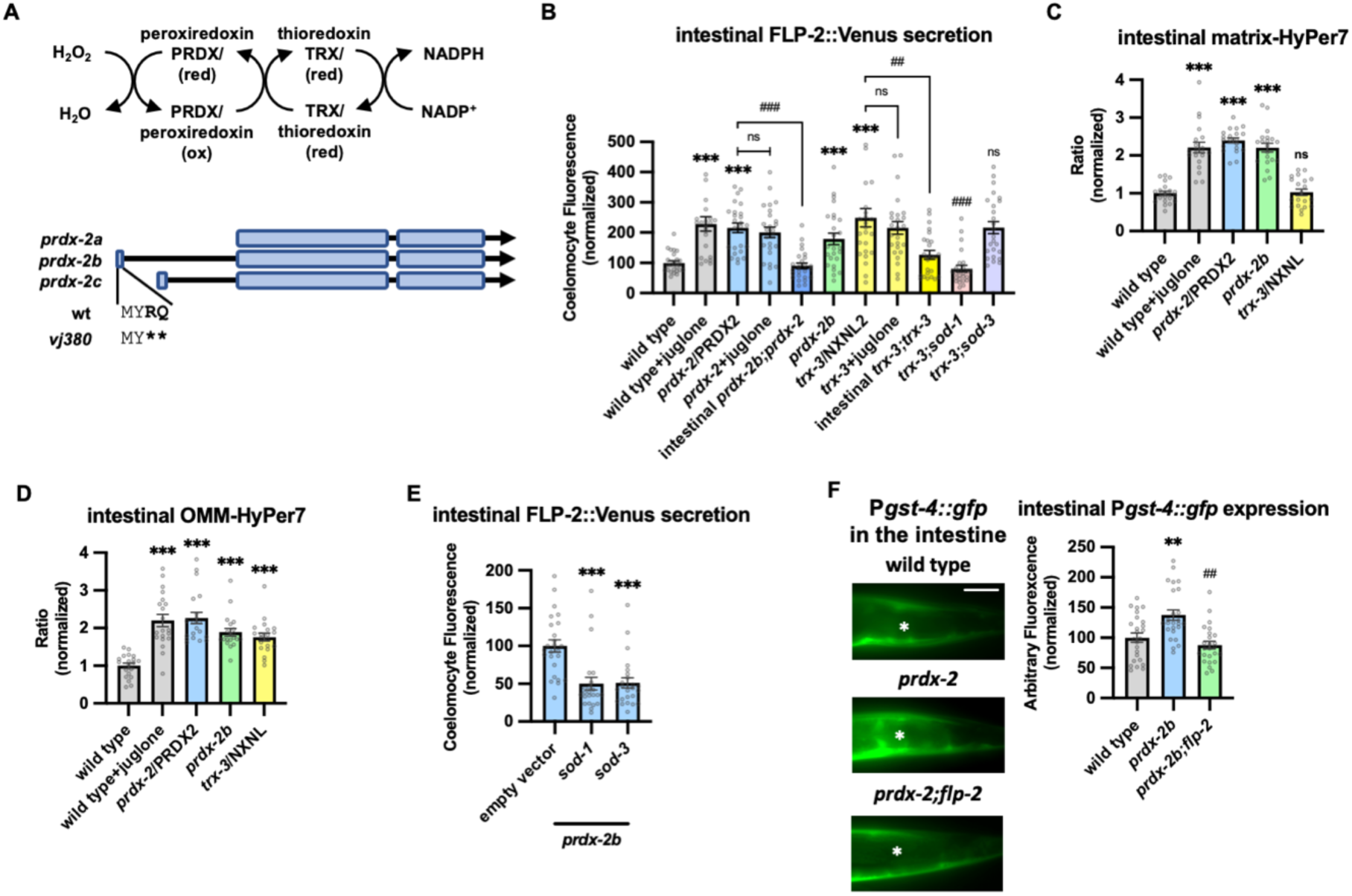
PRDX-2/PRDX and TRX-3/TRX regulate endogenous H_2_O_2_ and FLP-2 secretion. **A** (Top) Schematic showing the PRDX/TRX system in H_2_O_2_ detoxification. (Bottom) Schematic showing the three isoforms of *prdx-2* transcripts and *vj380* allele of *prdx-2b* knockout. **B** Quantification of average coelomocyte fluorescence of the indicated mutants expressing FLP-2::Venus fusion proteins in the intestine following M9 or 300μM juglone treatment for 10min. Intestinal *prdx-2b* denotes expression of *prdx-2b* cDNA under the *ges-1* promoter. Intestinal *trx-3* denotes expression of *trx-3* cDNA under the *ges-1* promoter. Unlined *** denotes statistical significance compared to “wild type”; unlined ## and ### denote statistical significance compared to “*trx-3”*. n= 25, 23, 25, 25, 25, 25, 25, 25, 25, 25, 25 independent animals. **C and D** Quantification of fluorescence in the posterior region of the indicated transgenic animals co-expressing matrix-HyPer7 (C) or OMM-HyPer7 (D) with TOMM-20::mCherry following M9 or 300μM juglone treatment. Ratio of images taken with 500nM (GFP) and 400nM (CFP) for excitation and 520nm for emission was used to measure H_2_O_2_ levels. Unlined *** and ns denote statistical significance compared to “wild type. (C) n = 20, 20, 20, 20, 20 independent animals. (D) n = 20, 20, 20, 20, 20 independent animals. **E** Quantification of average coelomocyte FLP-2::Venus fluorescence of transgenic animals fed with RNAi bacteria targeting the indicated genes following M9 treatment for 10min. Unlined *** denotes statistical significance compared to “empty vector”. n = 25, 23, 24 independent animals. **F** Representative images and quantification of average fluorescence in the posterior region of transgenic animals expressing P*gst-4::gfp* after 1h M9 or juglone exposure and 3h recovery. Asterisks mark the intestinal region for quantification. P*gst-4::gfp* expression in the body wall muscles, which appears as fluorescence on the edge animals in some images, was not quantified. Unlined ** denotes statistical significance compared to “wild type”, unlined ## denotes statistical analysis compared to “*prdx-2b”.* n = 25, 25, 25 independent animals. Scale bar: 10μM. **B-F** Data are mean values ± s.e.m normalized to wild type controls. n.s. not significant, ** and ## *P <* 0.01, *** and ### *P <* 0.001 by Brown-Forsythe and Welch ANOVA with Dunnett’s T3 multiple comparisons test.

There are three isoforms of *prdx-2* that arise by the use of alternative transcriptional start sites (Fig. 4A). Expressing the *prdx-2b* isoform selectively in the intestine fully rescued the elevated FLP-2::Venus secretion defects of *prdx-2* mutants, whereas expressing *prdx-2a* or *prdx-2c* isoforms failed to rescue (Fig. 4B, S4B and C). To independently verify the role of *prdx-2b* function in FLP-2 release, we generated a *prdx-2b*-specific knockout mutant by introducing an in frame stop codon within the *prdx-2b*-specific exon 1 using CRISPR/Cas9 (*prdx-2b(vj380)* Fig. 4A). *prdx-2b(vj380)* mutants exhibited increased H_2_O_2_ levels in the mitochondrial matrix and OMM (Fig. 4C and D), as well as increased FLP-2::Venus secretion compared to wild type controls that were indistinguishable from *prdx-2* null mutants (Fig. 4B). *prdx-2b* mutations could no longer increase FLP-2 secretion when either *sod-1* or *sod-3* activity was impaired (Fig. 4E and Fig. S4D). Thus, the *prdx-2b* isoform normally inhibits FLP-2 secretion likely by promoting the consumption of H_2_O_2_ in the mitochondrial matrix and/or cytosol.

Once oxidized, peroxiredoxins are reduced by thioredoxins (TRXs) for reuse (Netto and Antunes 2016). TRX-3 is an intestine-specific thioredoxin promoting protection against specific pathogen infections (Jiménez-Hidalgo et al. 2014; Miranda-Vizuete, Damdimopoulos, and Spyrou 2000; Netto and Antunes 2016). Mutations in *trx-3* elevated FLP-2::Venus release in the absence of juglone and expressing *trx-3* transgenes in the intestine restored wild type FLP-2 release to *trx-3* mutants (Fig. 4B). Juglone treatment failed to further enhance FLP-2::Venus release in *trx-3* mutants (Fig. 4B). Mutations in cytoplasmic *sod-1* but not in mitochondrial *sod-3* reduced the elevated FLP-2::Venus release in *trx-3* mutants to wild type levels (Fig. 4B). Mutations in *trx-3* increased H_2_O_2_ levels in the OMM but had no effect on matrix H_2_O_2_ levels (Fig. 4C and D). Thus TRX-3 likely functions in the cytosol but not in the matrix to neutralize H_2_O_2_, and elevated H_2_O_2_ levels in the cytosol are sufficient to drive FLP-2 secretion without SOD-3-mediated H_2_O_2_ generation in the matrix.

Finally, to investigate the physiological significance of elevated endogenous H_2_O_2_ levels on the oxidative stress response we examined the effects of *prdx-2b* mutations on expression of *gst-4*. *prdx-2b* mutants had significantly increased P*gst-4::gfp* expression in the intestine compared to wild type controls (Fig. 4F). The increased P*gst-4::gfp* expression in *prdx-2b* mutants was completely dependent upon *flp-2* signaling, since *gst-4* expression was reduced to wild type levels in *prdx-2b; flp-2* double mutants (Fig. 4F). Together our data suggest that *prdx-2b* functions in the intestine to maintain redox homeostasis following SOD-1/SOD-3 mediated H_2_O_2_ production by regulating the secretion of FLP-2 (Fig. 3I).

### PKC-2/PKCα/β mediates H_2_O_2_ induced FLP-2 secretion from the intestine

H_2_O_2_ functions as a cellular signaling molecule by oxidizing reactive cysteines to sulfenic acid, and this modification on target proteins can regulate intracellular signaling pathways (García Santamarina, Boronat i Llop, and Hidalgo Hernando 2014). One of the validated targets of H_2_O_2_ signaling is the protein kinase C (PKC) family of serine threonine kinases (Jia and Sieburth 2021; Konishi et al. 1997, 2001; Min, Kim, and Exton 1998). *C. elegans* encodes four PKC family members including *pkc-*1 and *pkc-*2, which are expressed at highest levels in the intestine (Islas-Trejo et al. 1997; Taylor et al. 2021). *pkc-1* null mutants had no effect on baseline or juglone-induced FLP-2 secretion (Fig. S5A). *pkc-2* null mutations did not alter baseline intestinal FLP-2 secretion, but they eliminated juglone-induced FLP-2 secretion (Fig. 5A). *pkc-2* encodes a calcium and diacylglycerol (DAG) stimulated PKCα/β protein kinase C that regulates thermosensory behavior by promoting transmitter secretion (Edwards et al. 2012; Land and Rubin 2017). Expressing *pkc-2* cDNA selectively in the intestine fully restored juglone-induced FLP-2 secretion to *pkc-2* mutants (Fig. 5A), whereas expressing a catalytically inactive *pkc-2(K375R)* variant (Van et al. 2021) failed to rescue (Fig. 5A). The intestinal site of action of *pkc-2* is in line with prior studies showing that *pkc-2* can function in the intestine to regulate thermosensory behavior (Land and Rubin 2017). *pkc-2* mutants exhibited wild type H_2_O_2_ levels in the mitochondrial matrix and OMM of the intestine in both the presence and absence of juglone (Fig. 5B and C). Increasing H_2_O_2_ levels by either acute H_2_O_2_ treatment or by *prdx-2* mutation failed to increase FLP-2 secretion in *pkc-2* mutants (Fig. 5D and E). To determine whether *pkc-2* can regulate the intestinal secretion of other peptides that are not associated with oxidative stress, we examined expulsion frequency, which is a measure of intestinal NLP-40 secretion (Mahoney et al. 2008; Wang et al. 2013). *pkc-2* mutants showed wild type expulsion frequency (Fig. S5B), indicating that intestinal NLP-40 release is largely unaffected. Together, these results show that *pkc-2* is not a general regulator of intestinal peptide secretion and instead functions downstream or in parallel to H_2_O_2_ to selectively promote FLP-2 secretion by a mechanism that involves phosphorylation of target proteins.

**Figure 5.**
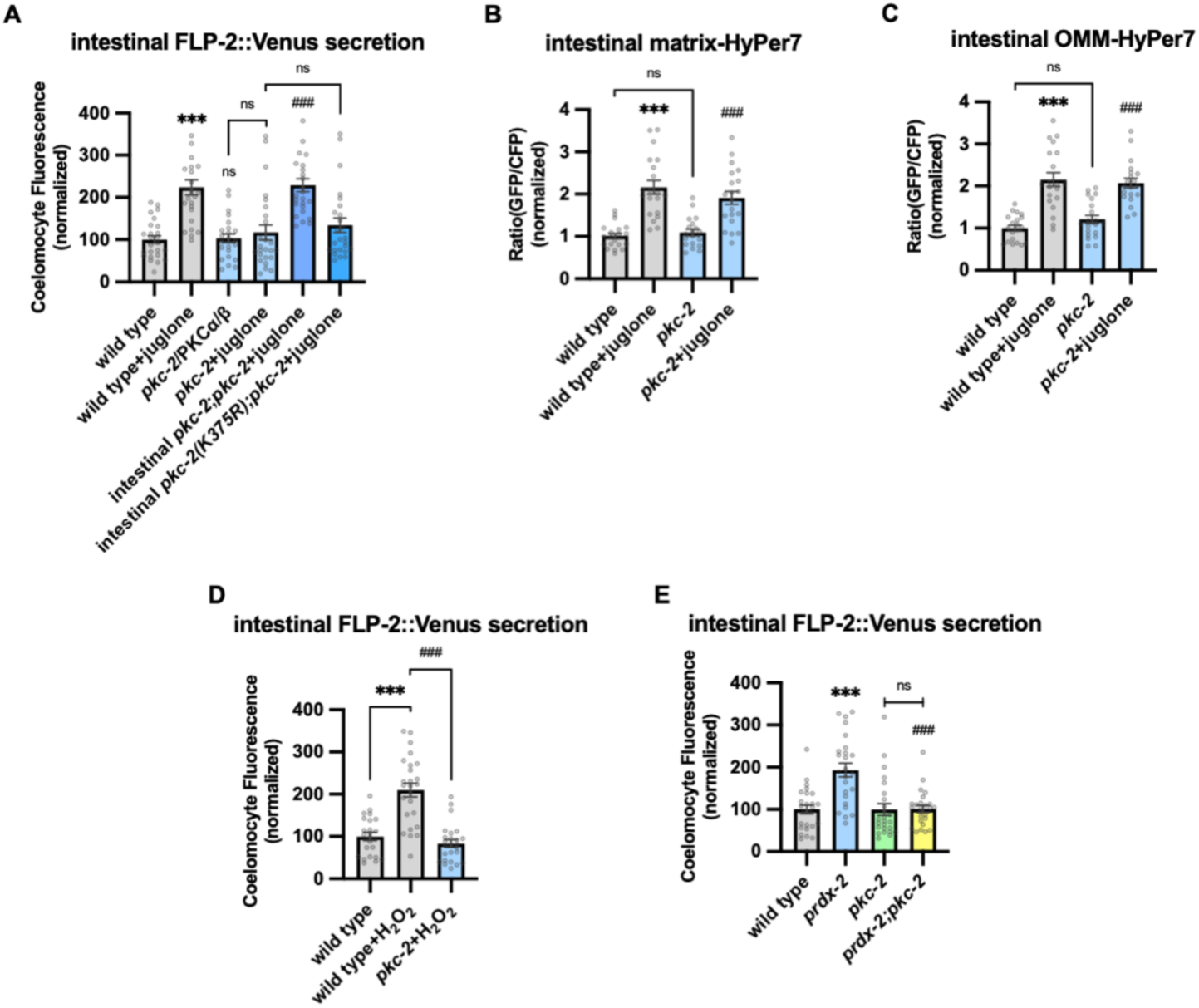
PKC-2/PKCα/β activation by H_2_O_2_ promotes FLP-2 secretion from the intestine. **A** Quantification of average coelomocyte fluorescence of the indicated mutants expressing FLP-2::Venus fusion proteins in the intestine following M9 or 300μM juglone treatment for 10min. Intestinal *pkc-2* denotes expression of *pkc-2b cDNA* under the ges-1 promoter. Intestinal *pkc-2b(K375R)* denotes expression of *pkc-2b(K375R)* variants under the *ges-1* promoter. Unlined *** and ns denote statistical significance compared to “wild type”; *###* denote statistical significance compared to “*pkc-*2+juglone”. n = 24, 24, 25, 25, 25, 25 independent animals. **B and C** Quantification of fluorescence in the posterior region of the indicated transgenic animals co-expressing matrix-HyPer7 (B) or OMM-HyPer7 (C) with TOMM-20::mCherry following M9 or 300μM juglone treatment. Ratio of images taken with 500nM (GFP) and 400nM (CFP) for excitation and 520nm for emission was used to measure H_2_O_2_ levels. Unlined *** denotes statistical significance compared to “wild type”; unlined ### denotes statistical analysis compared to “*pkc-2”.* (B) n = 20, 20, 19, 20 independent animals, (C) n = 20, 20, 20, 20 independent animals. **D** Quantification of average coelomocyte fluorescence of the indicated mutants expressing FLP-2::Venus fusion proteins in the intestine following M9 or 1mM H_2_O_2_ treatment for 10min. n = 23, 25, 25 independent animals. **E** Quantification of average coelomocyte fluorescence of the indicated mutants expressing FLP-2::Venus fusion proteins in the intestine following M9 treatment for 10min. Unlined *** denotes statistical significance compared to “wild type”; unlined ### denotes statistical significance compared to “*prdx-2”*. n = 25, 25, 25, 25 independent animals. **A-E** Data are mean values ± s.e.m normalized to wild type controls. ns. not significant, *** and *### P <* 0.001 by Brown-Forsythe and Welch ANOVA with Dunnett’s T3 multiple comparisons test.

### DAG positively regulates FLP-2 secretion

PKCα/β family members contain two N terminal C1 domains (C1A and C1B) whose binding to DAG promotes PKC recruitment to the plasma membrane (Burns and Bell 1991; Darby, Meng, and Fehrenbacher 2017; Johnson, Giorgione, and Newton 2000; Kim et al. 2016; Ono et al. 1989; Yanase et al. 2011). To address the role of DAG in promoting FLP-2 secretion by PKC-2, we examined mutants that are predicted to have altered DAG levels. Phosphatidylinositol phospholipase C beta (PLCβ) converts phosphatidyl inositol phosphate (PIP2) to DAG and inositol triphosphate (IP3, Fig. 6A), and impairing PLC activity leads to reduced cellular DAG levels (Nebigil 1997). *C. elegans* encodes two PLC family members whose expression is enriched in the intestine, *plc-2/* PLCβ and *egl-8/* PLCβ (Taylor et al. 2021). *plc-2* null mutants exhibited baseline and juglone-included FLP-2 secretion that were similar to wild type controls (Fig. S6). *egl-8* loss-of-function mutants exhibited wild type baseline FLP-2 secretion, but juglone-induced FLP-2 secretion was completely blocked (Fig. 6B). H_2_O_2_ levels in *egl-8* mutants were similar to wild type controls, both in the presence and absence of juglone (Fig. 6C and D). Thus, *egl-8/*PLCβ functions downstream of or in parallel to H_2_O_2_ production to promote FLP-2 secretion.

**Figure 6.**
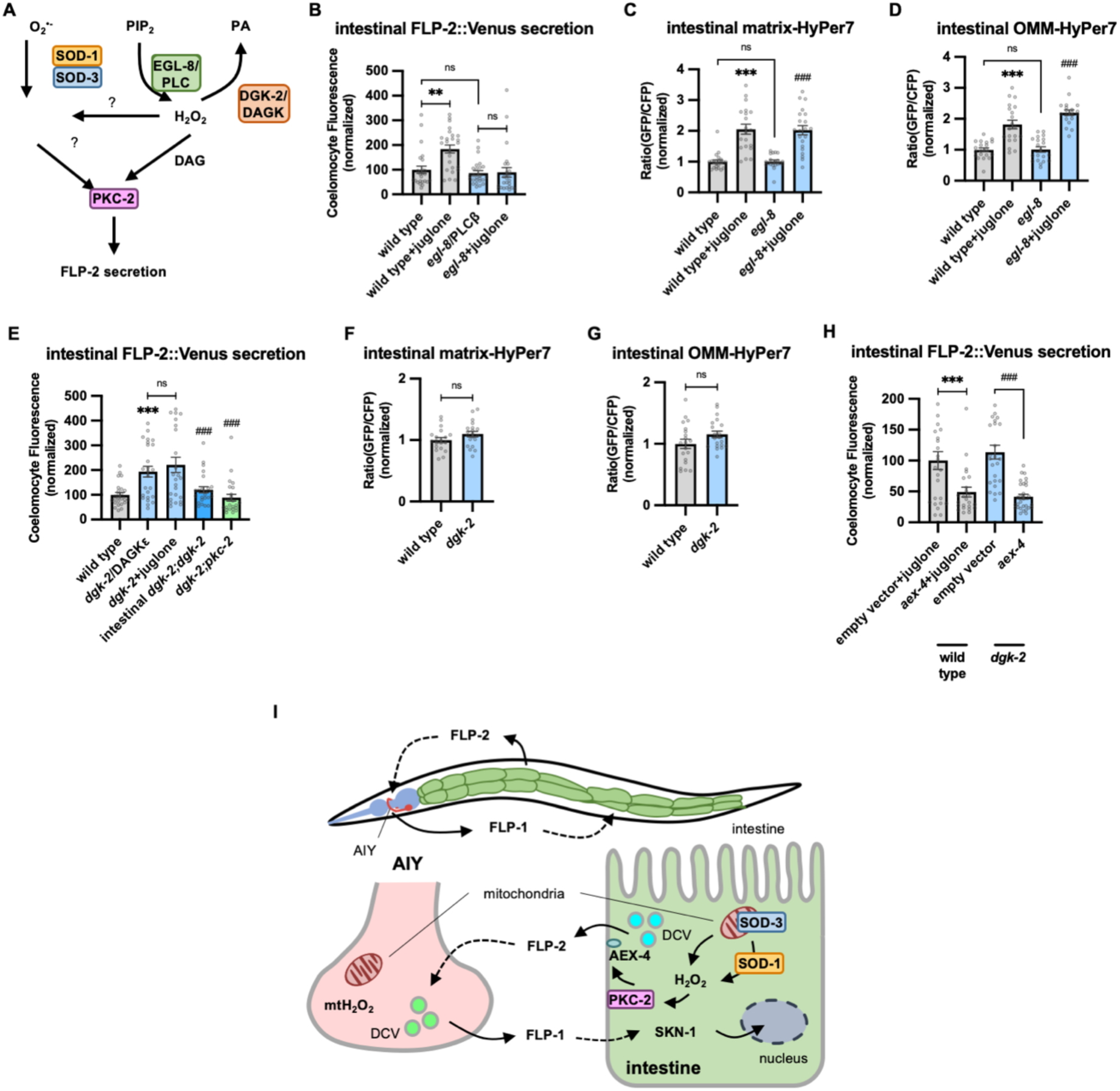
DAG promotes PKC-2 mediated FLP-2 secretion from the intestine. **A** Schematic showing PLC and DGK mediates DAG metabolism and DAG functions in H_2_O_2_-mediated FLP-2 signaling. **B** Quantification of average coelomocyte fluorescence of the indicated mutants expressing FLP-2::Venus fusion proteins in the intestine following M9 or juglone treatment for 10min. n = 25, 25, 25, 25 independent animals. **C and D** Quantification of fluorescence in the posterior region of the indicated transgenic animals co-expressing matrix-HyPer7 (C) or OMM-HyPer7 (D) with TOMM-20::mCherry following M9 or 300μM juglone treatment. Ratio of images taken with 500nM (GFP) and 400nM (CFP) for excitation and 520nm for emission was used to measure H_2_O_2_ levels. Unlined *** denotes statistical significance compared to “wild type”; unlined ### denotes statistical significance compared to “*egl-8”.* (C) n = 22, 20, 20, 21 independent animals, (D) n = 20, 20, 20, 20 independent animals. **E** Quantification of average coelomocyte fluorescence of the indicated mutants expressing FLP-2::Venus fusion proteins in the intestine following M9 or 300μM juglone treatment for 10min. Intestinal *dgk-2* denotes expression of *dgk-2a* cDNA under the *ges-1* promoter. Unlined *** denotes statistical significance compared to “wild type”; unlined ### denotes statistical significance compared to “*dgk-2*/DGKε”. n = 25, 25, 25, 25, 24 independent animals. **F and G** Quantification of fluorescence in the posterior region of the indicated transgenic animals co-expressing matrix-HyPer7 (F) or OMM-HyPer7 (G) with TOMM-20::mCherry following M9 treatment. Ratio of images taken with 500nM (GFP) and 400nM (CFP) for excitation and 520nm for emission was used to measure H_2_O_2_ levels. (F) n = 20, 20 independent animals, (G) n = 20, 20 independent animals. **H** Quantification of average coelomocyte fluorescence of the indicated transgenic animals fed with RNAi bacteria targeting the indicated genes in the intestine following M9 treatment for 10min. n = 25, 24, 25, 30 independent animals. **I** (Top) Schematic showing the position of intestine and AIY neurons in FLP-1-FLP-2 mediated axis. (Bottom) Schematic showing endogenous H_2_O_2_ promotes PKC-2/AEX-4 mediated FLP-2 release from the intestine in FLP-1-FLP-2 regulated inter-tissue axis. **B-H** Data are mean values ± s.e.m normalized to wild type controls. **B-E and H** ns. not significant, ** *P* < 0.01, *** and *### P <* 0.001 by Brown-Forsythe and Welch ANOVA with Dunnett’s T3 multiple comparisons test. **F and G** ns. not significant by unpaired t test with Welch’s correction.

DAG kinase converts DAG into phosphatidic acid (PA), and is therefore a negative regulator or DAG levels (Fig. 6A) (van Blitterswijk and Houssa 2000; Topham 2006). In *C. elegans*, *dgk-2*/DGKε is the highest expressing DAG kinase in the intestine. Mutations in *dgk-2* elevated FLP-2::Venus secretion (Fig. 6E) without altering H_2_O_2_ levels in the intestinal mitochondrial matrix or OMM (Fig. 6F and G). Expressing *dgk-2* transgenes selectively in the intestine restored normal FLP-2::Venus secretion to *dgk-2* mutants (Fig. 6E). Finally, the increase in FLP-2 secretion in *dgk-2* mutants was not further increased by juglone treatment, but it was completely blocked by *pkc-2* mutations or *aex-4*/SNAP25 mutations (Fig. 6E and H). These results show that FLP-2 secretion can be regulated bidirectionally by DAG, and they suggest that DAG and H_2_O_2_ function in a common genetic pathway upstream of *pkc-2* to promote FLP-2 secretion (Fig. 6A).

## Discussion

By screening for intercellular regulators of FLP-1 signaling from the nervous system in promoting the anti-oxidant response, we have uncovered a function for peptidergic signaling in mediating gut-to-neuron regulation of the anti-oxidant response in *C. elegans.* We identified the neuropeptide-like protein FLP-2 as an inter-tissue signal originating in the intestine to potentiate stress-induced FLP-1 release from AIY neurons and the subsequent activation of SKN-1 in the intestine. We found that H_2_O_2_ generated endogenously in the intestine or exogenously by acute oxidant exposure increases FLP-2 secretion from intestinal DCVs. H_2_O_2_ promotes FLP-2 exocytosis through PKC-2, and AEX-4/SNAP25. The use of oxidant-regulated peptide secretion exemplifies a mechanism that can allow the gut and the nervous system to efficiently and rapidly communicate through endocrine signaling to promote organism-wide protection in the face of intestinal stress (Fig. 6I).

### A new function for *flp-2* signaling in the antioxidant response

Previous studies have identified roles for *flp-2* signaling in development and in stress responses. *flp-2* promotes locomotion during molting (Chen et al. 2016) promotes entry into reproductive growth (Chai et al. 2022), regulates longevity (Kageyama et al. 2022), and activates the UPR^mt^ cell-non autonomously during mitochondrial stress (Shao et al. 2016). The function we identified for *flp-2* in the antioxidant response has some notable similarities with *flp-2*’s other functions. First, *flp-2* mediates its effects at least in part by regulating signaling by other peptides. *flp-2* signaling increases the secretion of the neuropeptide like protein PDF-1 during lethargus (Chen et al. 2016) and INS-35/insulin-like peptide for its roles in reproductive growth choice and longevity (Kageyama et al. 2022), in addition to regulating AIY FLP-1 secretion (Fig. 5G).. Second, the secretion of FLP-2 is dynamic. FLP-2 secretion decreases during lethargus (Chen et al. 2016) and increases under conditions that do not favor reproductive growth (Kageyama et al. 2022), as well increasing in response to oxidants (Fig. 2C and D). However, in some instances, the regulation of FLP-2 secretion may occur at the level of flp-2 expression (Kageyama et al. 2022), rather than at the level of exocytosis (Fig. 2C). Finally, genetic analysis of *flp-2* has revealed that under normal conditions, *flp-2* signaling may be relatively low, since *flp-2* mutants show no defects in reproductive growth choice when animals are well fed (Chai et al. 2022), show only mild defects in locomotion during molting in non-sensitized genetic backgrounds (Chen et al. 2016), and do not have altered baseline FLP-1 secretion or antioxidant gene expression in the absence of exogenous oxidants (Fig. 1D and E). It is notable that increased ROS levels are associated with molting (Back et al. 2012; Knoefler et al. 2012), ageing (Back et al. 2012; Van Raamsdonk and Hekimi 2010), starvation (Tao et al. 2017), and mitochondrial dysfunction (Dingley et al. 2010), raising the possibility that *flp-2* may be used in specific contexts associated with high ROS levels to affect global changes in physiology, behavior and development.

One major difference we found for *flp-2* signaling in our study is that intestinal, but not neuronal *flp-2* activates the oxidative stress response, whereas *flp-2* originates from neurons for its reported roles in development and the UPR^mt^. The intestine is uniquely poised to relay information about diet to the rest of the animal, and secretion of a number of neuropeptide-like proteins from the intestine (e.g, INS-11, PDF-2 and INS-7) is proposed to regulate responses to different bacterial food sources (Lee and Mylonakis 2017; Murphy, Lee, and Kenyon 2007; O’Donnell et al. 2018). Since bacterial diet can impact ROS levels in the intestine (Pang and Curran 2014), secretion of FLP-2 from the intestine could function to relay information about bacterial diet to distal tissues to regulate redox homeostasis. In addition, the regulation of intestinal FLP-2 release by oxidants may meet a unique spatial, temporal or concentration requirement for activating the antioxidant response that cannot be met by its release from the nervous system.

### AIY as a target for *flp-2* signaling

AIY interneurons receive sensory information from several neurons primarily as glutamatergic inputs to regulate behavior (Bargmann et al. 2007; Clark et al. 2006; Satoh et al. 2014). Our study reveals a previously undescribed mechanism by which AIY is activated through endocrine signaling originating from FLP-2 secretion from the intestine. FLP-2 could act directly on AIY, or it may function indirectly through upstream neurons that relay FLP-2 signals to AIY. *frpr-18* encodes an orexin-like GPCR that can be activated by FLP-2-derived peptides in transfected mammalian cells (Larsen et al. 2013; Mertens et al. 2005), and *frpr-18* functions downstream of *flp-2* in the locomotion arousal circuit (Chen et al. 2016). *frpr-18* is expressed broadly in the nervous system including in AIY (Chen et al. 2016), and loss-of-function *frpr-18* mutations lead to hypersensitivity to certain oxidants (Ouaakki et al. 2023). FRPR-18 is coupled to the heterotrimeric G protein Gαq (Larsen et al. 2013; Mertens et al. 2005), raising the possibility that FLP-2 may promote FLP-1 secretion from AIY by directly activating FRPR-18 in AIY. However, *flp-2* functions independently of *frpr-18* in the reproductive growth circuit, and instead functions in a genetic pathway with the GPCR *npr-30* (Chai et al. 2022). In addition, FLP-2-derived peptides (of which there are at least three) can bind to the GPCRs DMSR-1, or FRPR-8 in transfected cells (Beets et al. 2023). Identifying the relevant FLP-2 peptide(s), the FLP-2 receptor and its site of action will help to define the circuit used by intestinal *flp-2* to promote FLP-1 release from AIY.

FLP-1 release from AIY is positively regulated by H_2_O_2_ generated from mitochondria (Jia and Sieburth 2021). Here we showed that H_2_O_2_-induced FLP-1 release requires intestinal *flp-2* signaling. However, *flp-2* does not appear to promote FLP-1 secretion by increasing H_2_O_2_ levels in AIY (Fig 1E), and *flp-2* signaling is not sufficient to promote FLP-1 secretion in the absence of H_2_O_2_ (Fig. 1D). These results point to a model whereby at least two conditions must be met in order for AIY to increase FLP-1 secretion: an increase in H_2_O_2_ levels in AIY itself, and an increase in *flp-2* signaling from the intestine. Thus AIY integrates stress signals from both the nervous system and the intestine to activate the intestinal antioxidant response through FLP-1 secretion. The requirement of signals from multiple tissues for FLP-1 secretion may function to limit the activation of SKN-1, since unregulated SKN-1 activation can be detrimental to organismal health (Turner, Ramos, and Curran 2024). AIY shows a sporadic Ca^2+^ response regardless of the presence of explicit stimulation (Ashida, Hotta, and Oka 2019; Bargmann et al. 2007; Clark et al. 2006), and FLP-1 secretion from AIY is calcium dependent (Jia and Sieburth 2021)How mitochondrial H_2_O_2_ levels are established in AIY by intrinsic or extracellular inputs, and how AIY integrates H_2_O_2_ and *flp-2* signaling to control FLP-1 secretion remain to be defined.

### A role for endogenous H_2_O_2_ in regulated neuropeptide secretion

Using HyPer7, we showed that acute juglone exposure results in a rapid elevation of endogenous H_2_O_2_ levels inside and outside intestinal mitochondria and a corresponding increase of FLP-2 release from the intestine that depends on the cytoplasmic superoxide dismutase *sod-1*, and mitochondrial *sod-3*. We favor a model whereby superoxide generated by juglone in the mitochondria is converted to H_2_O_2_ by SOD-3 in the matrix and by SOD-1 in the cytosol. In this case, both the superoxide generated by juglone and the H_2_O_2_ generated by SOD-3 would have to be able to exit the mitochondria and enter the cytosol. Superoxide and H_2_O_2_ can be transported across mitochondrial membranes through anion channels and aquaporin channels, respectively (Bienert and Chaumont 2014; Ferri et al. 2003; Han et al. 2003; Kontos et al. 1985). The observation that both SOD-1 and SOD-3 activity are necessary to drive FLP-2 release suggests that H_2_O_2_ levels much reach a certain threshold in the cytoplasm to promote FLP-2 release, and this threshold requires the generation of H_2_O_2_ by both SOD-1 and SOD-3.

We identified a role for the antioxidant peroxiredoxin-thioredoxin system, encoded by *prdx-2* and *trx-3*, in maintaining low endogenous H_2_O_2_ levels in the intestine and in negatively regulating FLP-2 secretion. We showed that the *prdx-2b* isoform functions to inhibit FLP-2 secretion and to lower H_2_O_2_ levels in both the mitochondrial matrix and on the cytosolic side of mitochondria. These observations are consistent with a subcellular site of action for PRDX-2B in either the matrix only or in both the matrix and cytosol. In contrast, *trx-3* mutations do not alter mitochondrial H_2_O_2_ levels, suggesting that TRX-3 functions exclusively in the cytosol. Thus, the PRDX-2B-TRX-3 combination may function in the cytosol, and PRDX-2B may function with a different TRX family member in the matrix. There are several thioredoxin-domain containing proteins in addition to *trx-3* in the *C. elegans* genome (including *trx-5*/NXNL2) that could be candidates for this role. Alternatively, *prdx-2* may function alone or with other redox proteins. PRDX-2 may function without thioredoxins in its roles in light sensing and stress response in worms (Li et al. 2016; Oláhová et al. 2008; Oláhová and Veal 2015). PRDX-2B contains a unique N terminal domain that is distinct from the catalytic domain and is not found on the other PRDX-2 isoforms. This domain may be important for targeting PRDX-2B to specific subcellular location(s) where it can regulate FLP-2 secretion.

### Regulation of FLP-2 exocytosis by PKC-2/PKCα/β and AEX-4/SNAP25

We demonstrated that *pkc-2* mediates the effects of H_2_O_2_ on intestinal FLP-2 secretion, and H_2_O_2_- and DAG-mediated PKC-2 activation are likely to function in a common genetic pathway to promote FLP-2 secretion. Our observations that DAG is required for the effects of juglone (Fig. 6B), are consistent with a two-step activation model for PKC-2, in which H_2_O_2_ could first modify PKC-2 in the cytosol, facilitating subsequent PKC-2 recruitment to the membrane by DAG. Alternatively, DAG could first recruit PKC-2 to membranes, where it is then modified by H_2_O_2_. We favor a model whereby H_2_O_2_ modification occurs in the cytosol, since H_2_O_2_ produced locally by mitochondria would have access to cytosolic pools of PKC-2 prior to its membrane translocation.

We defined a role for *aex-4*/SNAP25 in the fusion step of FLP-2 containing DCVs from the intestine under normal conditions as well as during oxidative stress. In neuroendocrine cells, phosphorylation of SNAP25 on Ser187 potentiates DCV recruitment into releasable pools (Nagy et al. 2002; Shu et al. 2008; Yang et al. 2007), and exocytosis stimulated by the DAG analog phorbol ester (Gao et al. 2016; Shu et al. 2008), without altering baseline SNAP25 function. Interestingly, the residue corresponding to Ser187 is conserved in AEX-4, raising the possibility that PKC-2 potentiates FLP-2 secretion by phosphorylating AEX-4. Since SNAP25 phosphorylation on Ser187 has been shown to increase its interaction with syntaxin and promote SNARE complex assembly in vitro (Gao et al. 2016; Yang et al. 2007), it is possible that elevated H_2_O_2_ levels could promote FLP-2 secretion by positively regulating SNARE-mediated DCV fusion at intestinal release sites on the basolateral membrane through AEX-4/SNAP25 phosphorylation by PKC-2. Prior studies have shown that PKC-2 phosphorylates the SNARE-associated protein UNC-18 in neurons to regulate thermosensory behavior (Edwards et al. 2012; Land and Rubin 2017). Thus, PKC-2 may have multiple targets in vivo and target selection may be dictated by cell type and/or the redox status of the cell.

### Similar molecular mechanisms regulating FLP-1 and FLP-2 release

The molecular mechanisms we identified that regulate FLP-2 secretion from the intestine are similar in several respects to those regulating FLP-1 secretion from AIY. First, the secretion of both peptides is positively regulated by H_2_O_2_ originating from mitochondria. Second, in both cases, H_2_O_2_ promotes exocytosis of neuropeptide-containing DCVs by a mechanism that depends upon the kinase activity of protein kinase C. Finally, the secretion of both peptides is controlled through the regulation of H_2_O_2_ levels by superoxide dismutases and by the peroxiredoxin-thioredoxin system. H_2_O_2_-regulated FLP-1 and FLP-2 secretion differ in the identity of the family members of some of the genes involved. *prdx-3-trx-2* and *sod-2* family members regulate H_2_O_2_ levels in AIY, whereas *prdx-2-trx-3* and *sod-1/sod-3* family members regulate H_2_O_2_ levels in the intestine. In addition, *pkc-1* promotes H_2_O_2_ induced FLP-1 secretion from AIY whereas *pkc-2* promotes H_2_O_2_-induced FLP-2 secretion from the intestine. Nonetheless, it is noteworthy that two different cell types utilize largely similar pathways for the H_2_O_2_-mediated regulation of neuropeptide release, raising the possibility that similar mechanisms may be utilized in other cell types and/or organisms to regulate DCV secretion in response to oxidative stress.

## Materials and Methods

### Strains and transgenic lines

*C. elegans* strains were maintained at 20°C in the dark on standard nematode growth medium (NGM) plates seeded with OP50 Escherichia coli as food source, unless otherwise indicated. All strains were synchronized by picking mid L4 stage animal either immediately before treatment (for coelomocyte imaging and intestine imaging) or 24h before treatment (for P*gst-4::gfp* imaging). The wild type strain was Bristol N2. Mutants used in this study were out-crossed at least four times.

Transgenic lines were generated by microinjecting plasmid mixes into the gonads of young adult animals following standard techniques (Mello et al. 1991). Microinjection mixes were prepared by mixing expression constructs with the co-injection markers pJQ70 (P*ofm-1::rfp*, 25ng/μL), pMH163 (P*odr-1::mCherry*, 40ng/μL), pMH164 (P*odr-1::gfp*, 40ng/μL) or pDS806 (P*myo-3::mCherry*, 20ng/μL) to a final concentration of 100ng/μL. For tissue-specific expression, a 1.5kb *rab-3* promoter was used for pan-neuronal expression (Nonet et al. 1997), a 2.0kb *ges-1* or a 3.5kb *nlp-40* promoter was used for intestinal expression (Egan et al. 1995; Wang et al. 2013). At least three transgenic lines were examined for each transgene, and one representative line was used for quantification. Strains and transgenic lines used in this study are listed in the Supplementary Table.

### Molecular Biology

All gene expression vectors were constructed with the backbone of pPD49.26. Promoter fragments including P*rab-3*, and P*ges-1* were amplified from genomic DNA; genes of interest, including cDNA fragments (*aex-5, snt-5, sod-1b, sod-3, isp-1, prdx-2a, prdx-2b, prdx-2c, trx-3, pkc-2b, dgk-2a, aex-4*) and genomic fragments (*flp-2, flp-40, nlp-36, nlp-27*) were amplified from cDNA library and genomic DNA respectively using standard molecular biology protocols. Expression plasmid of HyPer7 was designed based on reported mammalian expression plasmid for HyPer7 (Pak et al. 2020) and was synthesized by Thermo Fisher Scientific with codon optimization for gene expression in *C. elegans.* Plasmids and primers used in this study are listed in the Supplementary Table.

### Toxicity Assay

A stock solution of 50mM juglone in DMSO was freshly made on the same day of liquid toxicity assay. 120μM working solution of juglone in M9 buffer was prepared using stock solution before treatment. Around 60-80 synchronized adult animals were transferred into a 1.5mL Eppendorf tube with fresh M9 buffer and washed three times, and a final wash was done with either the working solution of juglone with or M9 DMSO at the concentrations present in juglone-treated animals does not contribute to toxicity since DMSO treatment alone caused no significant change in survival compared to M9-treated controls (Fig. S1C). Animals were incubated in the dark for 4h on rotating mixer before being transferred onto fresh NGM plates seeded with OP50 to recover in the dark at 20°C for 16 hours. Percentage of survival was assayed by counting the number of alive and dead animals. Toxicity assays were performed in triplicates.

### RNAi Interference

Plates for feeding RNAi interference were prepare as described (Kamath 2003). Around 20-25 gravid adult animals with indicated genotype were transferred onto the RNAi plates that were seeded with HT115(DE3) bacteria transformed with L4440 vectors with targeted gene inserts or empty L4440 vectors. Eggs were collected for 4h to obtain synchronized populations. L4 stage animals were collected for further assays. RNAi clones were from Ahringer or Vidal RNAi library, or made from genomic DNA. Details were listed in the Supplementary Table.

### Behavioral assays

The defecation motor program was assayed as previously described (Liu and Thomas 1994). Twenty to thirty L4 animals were transferred onto a fresh NGM plate seeded with OP50 *E. coli* and were stored in a 20°C incubator for 24 hours. After 24 hours, ten consecutive defecation cycles were observed from three independent animals and the mean and the standard error was calculated for each genotype. The pBoc and aBoc steps were recorded using custom Etho software (James Thomas Lab website:http://depts.washington.edu/jtlab/software/otherSoftware.html)

### Microscopy and Fluorescence Imaging

Approximately 30-40 age matched animals were paralyzed with 30mg/mL 2,3-butanedione monoxime (BDM) in M9 buffer and mounted on 2% agarose pads. Images were captured using the Nikon eclipse 90i microscope equipped with Nikon Plan Apo 20x, 40x, 60x, and 100x oil objective (N.A.=1.40), and a Photometrics Coolsnap ES2 camera or a Hamamatsu Orca Flash LT+CMOS camera. Metamorph 7.0 software (Universal Imaging/Molecular Devices) was used to capture serial image stacks and to obtain the maximum intensity projection image for analysis.

For transcriptional reporter imaging, young adult animals were transferred into a 1.5mL Eppendorf tube with M9 buffer, washed three times and incubated in 50μM working solution of juglone or M9 buffer control with equivalent DMSO for 1h in the dark on rotating mixer before recovering on fresh NGM plates with OP50 for 3h in the dark at 20°C. The posterior end of the intestine was imaged with the 60x objective and quantification for average fluorescence intensity of a 16-pixel diameter circle in the posterior intestine was calculated using Metamorph.

For coelomocyte imaging, L4 stage animals were transferred in fresh M9 buffer on a cover slide, washed six times with M9 before being exposed to 300μM juglone in M9 buffer (diluted from freshly made 50mM stock solution), 1mM H_2_O_2_ in M9 buffer, or M9 buffer. DMSO at the concentrations present in juglone-treated animals does not alter neuropeptide secretion since DMSO treatment alone caused no significant change in FLP-1::Venus or FLP-2::Venus coelomocyte fluorescence compared to M9-treated controls. (Fig. S1D and S2E). Animals were then paralyzed in BDM and images of coelomocytes next to the posterior end of intestine were taken using the 100x oil objective. Average fluorescence intensity of Venus from the endocytic compartments in the posterior coelomocytes was measured in ImageJ.

For fusion protein fluorescence imaging, L4 stage animals were exposed to M9 buffer or indicated oxidants for 10min before being paralyzed in BDM and images taken of the posterior end of the intestine using 100x oil objective. For HyPer7 imaging, Z stacks were obtained using GFP (excitation/emission: 500nm/520nm) and CFP (excitation/emission: 400nm/520nm) filter sets sequentially, HyPer7 fluorescence signal was quantified as the ratio of GFP to CFP fluorescence intensity changes with respect to the baseline [(Ft − F0)/F0].

### CRISPR/Cas9 Editing

*prdx-2b(vj380)* knock-out mutants were generated using a co-CRISPR protocol (Arribere et al. 2014). A sgRNA and a repair single stranded oligodeoxynucleotides (ssODN) targeting *dpy-10* were co-injected with a sgRNA for genes of interest and a ssODN that induces homology-directed repair (HDR) to introduce Cas9 mediated mutagenesis. Fifteen young adult animals were injected to produce around thirty singled F1 animals carrying Dpy or Rol phenotype. F2 animals were genotyped for mutations based on PCR and enzyme digest. Homozygous mutants were outcrossed with wild type animals at least four times before being used for assays.

### Statistics

Statistical analysis was performed on GraphPad Prism 9. Unpaired *t*-test with two tails was used for two groups and one-way ANOVA with multiple comparison corrections was used to three or more groups to determine the statistical significance. Statistical details and n are specified in the figure legends. All comparisons are conducted based on wild-type controls unless indicated by lines between genotypes. Bar graphs with plots were generated using GraphPad Prism 9.

## Acknowledgements

*C. elegans* strains used in this study were provided by the Caenorhabditis Genetics Centre (CGC), which is funded by the NIH National Center for Research Resources (NCRR). We thank members of the Sieburth lab for critical reading and discussion of the manuscript. This work was supported by grants from National Institute of Health NINDS R01NS071085 and R01NS110730 to D.S.

## Supplementary Figures

**Supplementary Figure 1.**
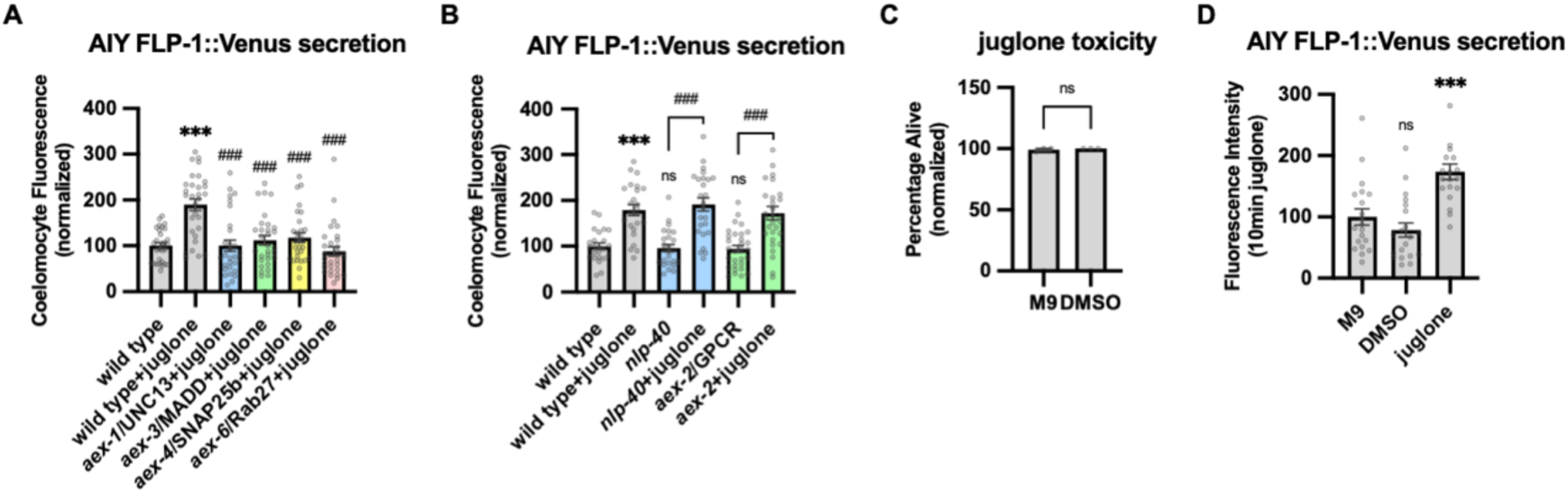
The effect of intestinal DCV secretion mutations on FLP-1 release from AIY. **A** Quantification of average coelomocyte fluorescence of the indicated mutants expressing FLP-1::Venus fusion proteins in AIY following M9 or 300μM juglone treatment for 10min. Unlined *** and *###* denotes statistical significance compared to “wild type”. n = 30, 30, 29, 30, 30, 30 independent animals. **B** Quantification of average coelomocyte fluorescence of the indicated mutants expressing FLP-1::Venus fusion proteins in AIY following M9 or 300μM juglone treatment for 10min. Unlined *** and ns denote statistical analysis compared to “wild type”. n = 24, 24, 25, 25, 30, 30 independent animals. **C** Average percentage of surviving young adult animals of the indicated genotypes after 16h recovery following 4h DMSO treatment. n = 203, 174 independent biological samples over three independent experiments. **D** Quantification of average coelomocyte fluorescence of the indicated mutants expressing FLP-1::Venus fusion proteins in AIY following M9, DMSO or juglone treatment for 10min. Unlined ns and *** denote statistical significance compared to “M9”. n = 20, 20, 19 independent animals. **A-D** Data are mean values ± s.e.m normalized to wild type controls. **A, B and D** ns. not significant, *** and ### *P* < 0.001 by Brown-Forsythe and Welch ANOVA with Dunnett’s T3 multiple comparisons test. **C** ns. not significant by unpaired t test with Welch’s correction.

**Supplementary Figure 2.**
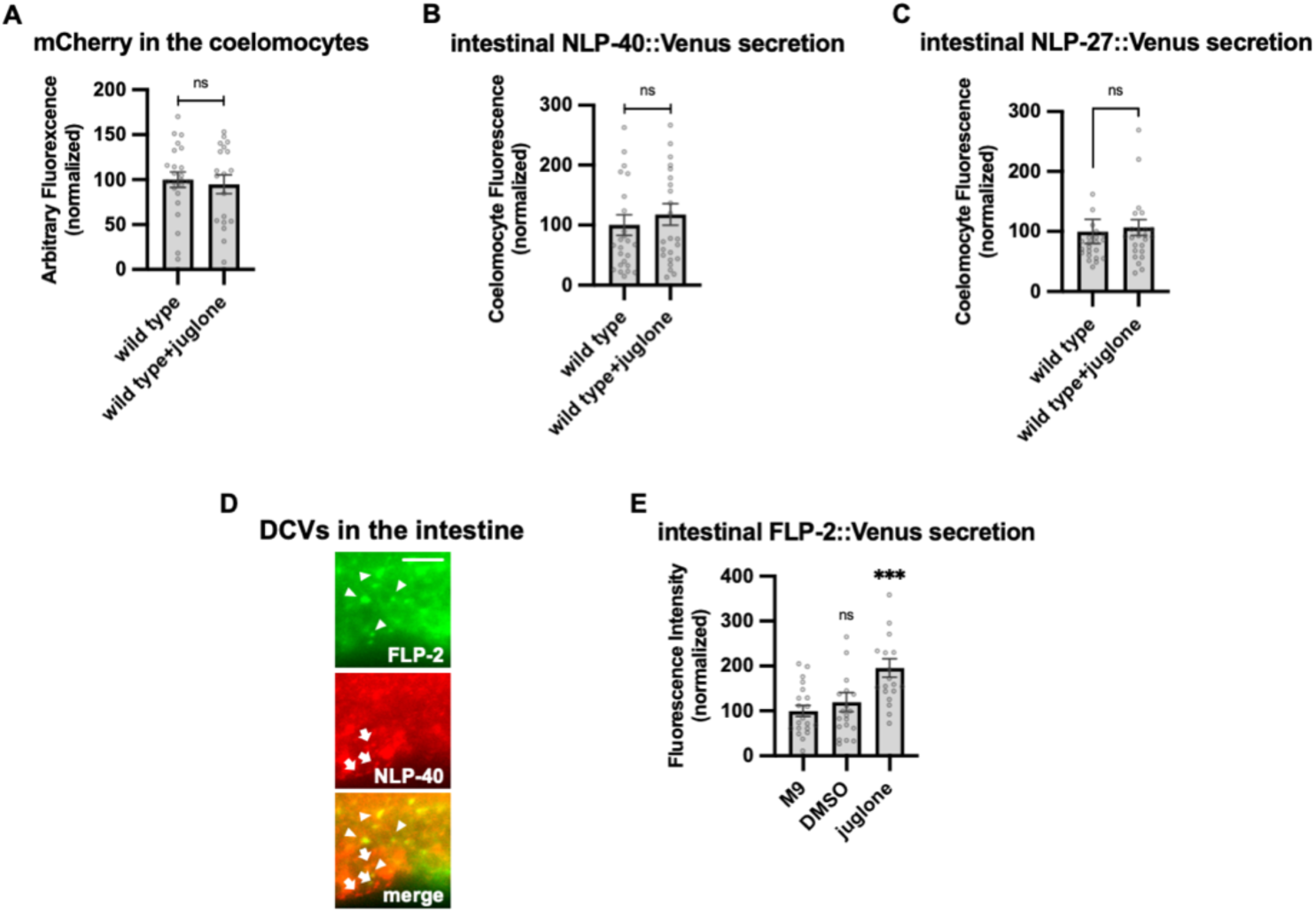
Specificity of juglone on intestinal peptide secretion, and FLP-2 and NLP-40 localization in the intestine. **A** Quantification of average coelomocyte fluorescence of the indicated mutants co-expressing FLP-2::Venus in the intestine (under the *ges-1* promoter) and mCherry in the coelomocytes (under the *ofm-1* promoter) following M9 or 300μM juglone treatment for 10min. n = 23, 19 independent animals. **B** Quantification of average coelomocyte fluorescence of transgenic animals expressing NLP-40::Venus fusion proteins in the intestine following M9 or 300μM juglone exposure for 10min. n = 25, 24 independent animals. **C** Quantification of average coelomocyte fluorescence of transgenic animals expressing NLP-27::Venus fusion proteins in the intestine following M9 or 300μM juglone exposure for 10min. n = 23, 25 independent animals. **D** Representative images of fluorescence distribution in the posterior intestinal region of transgenic animals co-expressing FLP-2::Venus fusion proteins (marked by arrowheads) and NLP-40::mTur2 fusion proteins (marked by arrows). Scale bar: 5μM. **E** Quantification of average coelomocyte fluorescence of transgenic animals expressing FLP-2::Venus fusion proteins in the intestine following M9, DMSO or 300μM juglone exposure for 10min. Unlined ns and *** denote statistical significance compared to “M9”. n = 20, 20, 20 independent animals. **A-C and E** Data are mean values ± s.e.m normalized to wild type controls. **A-C** ns. not significant by unpaired t test with Welch’s correction. **E** ns. not significant by Brown-Forsythe and Welch ANOVA with Dunnett’s T3 multiple comparisons test.

**Supplementary Figure 3.**
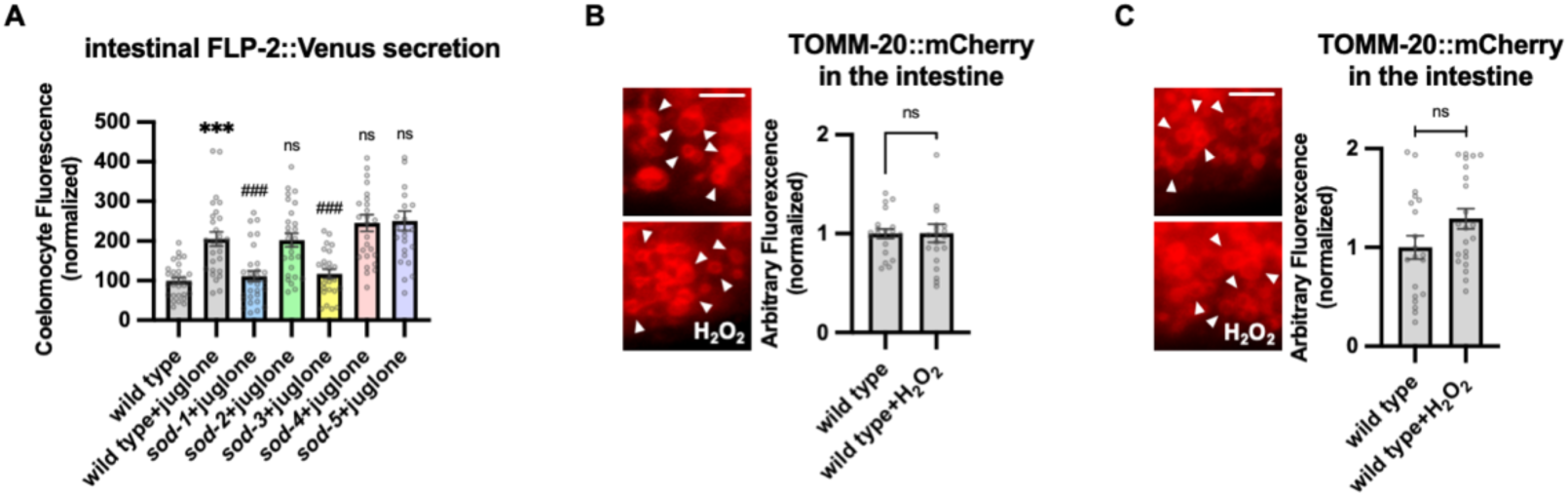
SODs function in juglone induced FLP-2 release from the intestine and mitochondrial mCherry control. **A** Quantification of average coelomocyte fluorescence of the indicated mutants expressing FLP-2::Venus fusion proteins in the intestine following M9 or juglone treatment for 10min. Unlined *** denotes statistical significance compared to “wild type”; unlined ### and ns denote statistical significance compared to “wild type + juglone” n = 29, 27, 29, 27, 25, 26, 24 independent animals. **B and C** Representative images and quantification of average fluorescence intensity of TOMM-20::mCherry proteins in transgenic animals co-expressing matrix-HyPer7 (B) or OMM-HyPer7 (C) following M9 or H_2_O_2_ treatment for 10min. (B) Scale bar: Scale bar: 5μM. n = 20, 20 independent animals. (C) Scale bar: Scale bar: 5μM. n = 20, 22 independent animals. **A-C** Data are mean values ± s.e.m normalized to wild type controls. **A** ns. not significant, *** and ### *P <* 0.001 by Brown-Forsythe and Welch ANOVA with Dunnett’s T3 multiple comparisons test. **B and C** ns. not significant by unpaired t test with Welch’s correction.

**Supplementary Figure 4.**
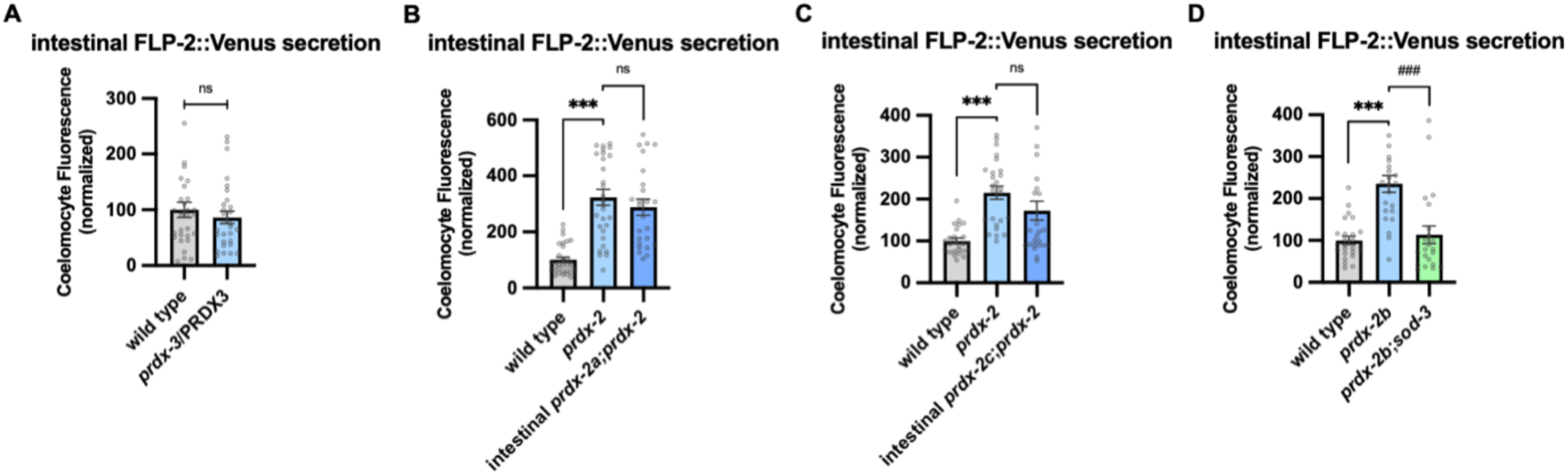
PRDX-2 intestinal rescue and mediates SOD-3 dependent regulation of FLP-2 release. **A** Quantification of average coelomocyte fluorescence of the indicated mutants expressing FLP-2::Venus fusion proteins in the intestine following M9 treatment for 10min. n = 30, 29 independent animals. **B** Quantification of average coelomocyte fluorescence of the indicated mutants expressing FLP-2::Venus fusion proteins in the intestine following M9 treatment for 10min. Intestinal *prdx-2a* denotes expression of *prdx-2a* cDNA under the *ges-1* promoter. n = 30, 30, 25 independent animals. **C** Quantification of average coelomocyte fluorescence of the indicated mutants expressing FLP-2::Venus fusion proteins in the intestine following M9 treatment for 10min. Intestinal *prdx-2c* denotes expression of *prdx-2c* cDNA under the *ges-1* promoter. n = 25, 25, 25 independent animals. **D** Quantification of average coelomocyte fluorescence of the indicated mutants expressing FLP-2::Venus fusion proteins in the intestine following M9 treatment for 10min. n = 25, 23, 22 independent animals. **A-D** Data are mean values ± s.e.m normalized to wild type controls. **A** ns. not significant by unpaired t test with Welch’s correction. **B-D** ns. not significant, *** and *### P <* 0.001 by Brown-Forsythe and Welch ANOVA with Dunnett’s T3 multiple comparisons test.

**Supplementary Figure 5.**
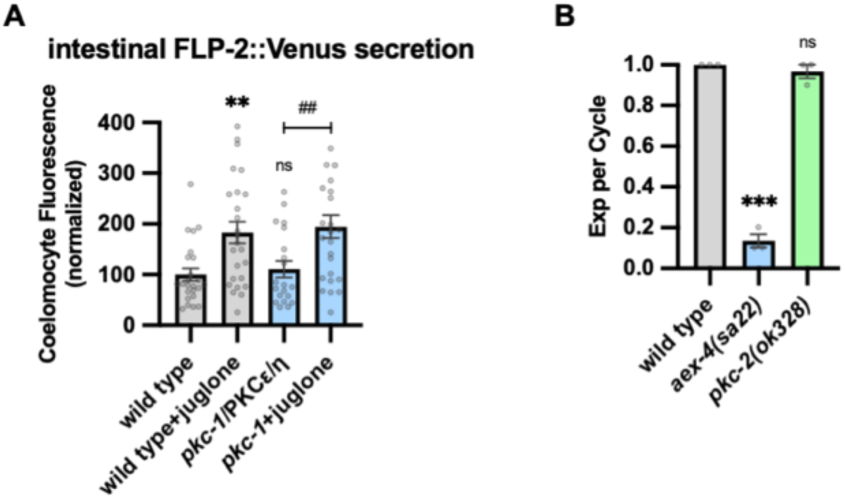
Juglone promotes FLP-2 release in *pkc-1* mutants and expulsion analysis. **A** Quantification of average coelomocyte fluorescence of the indicated mutants expressing FLP-2::Venus fusion proteins in the intestine following M9 or juglone treatment for 10min. Unlined ns and ** denote statistical significance compared to “wild type”. n = 24, 25, 20, 25 independent animals. **B** Quantification of the number of expulsions (Exp) per defecation cycle in adult animals of the indicated genotypes. Unlined *** and ns denote statistical significance compared to “wild type”. n = 30, 30, 30, 30 in three independent animals. **A-B** Data are mean values ± s.e.m normalized to wild type controls. ns. not significant, ** and ## *P* < 0.01, *** *P <* 0.001 by Brown-Forsythe and Welch ANOVA with Dunnett’s T3 multiple comparisons test.

**Supplementary Figure 6.**
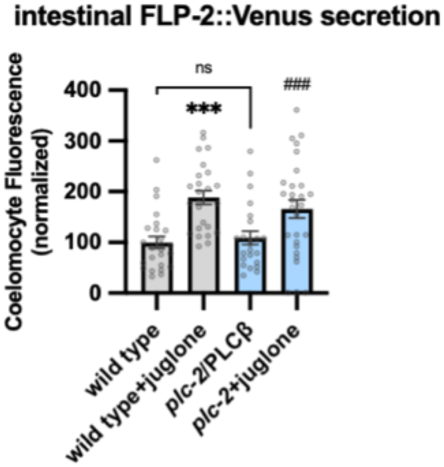
Juglone promotes FLP-2 release in *plc-2* mutants. Quantification of average coelomocyte fluorescence of the indicated mutants expressing FLP-2::Venus fusion proteins in the intestine following M9 or juglone treatment for 10min. Unlined *** denotes statistical significance compared to “wild type”; unlined ### denotes statistical significance compared to “*plc-2*/PLCβ”. n = 25, 25, 23, 28 independent animals. Data are mean values ± s.e.m normalized to wild type controls. ns. not significant, *** and *### P <* 0.001 by Brown-Forsythe and Welch ANOVA with Dunnett’s T3 multiple comparisons test.

**Table S1.**
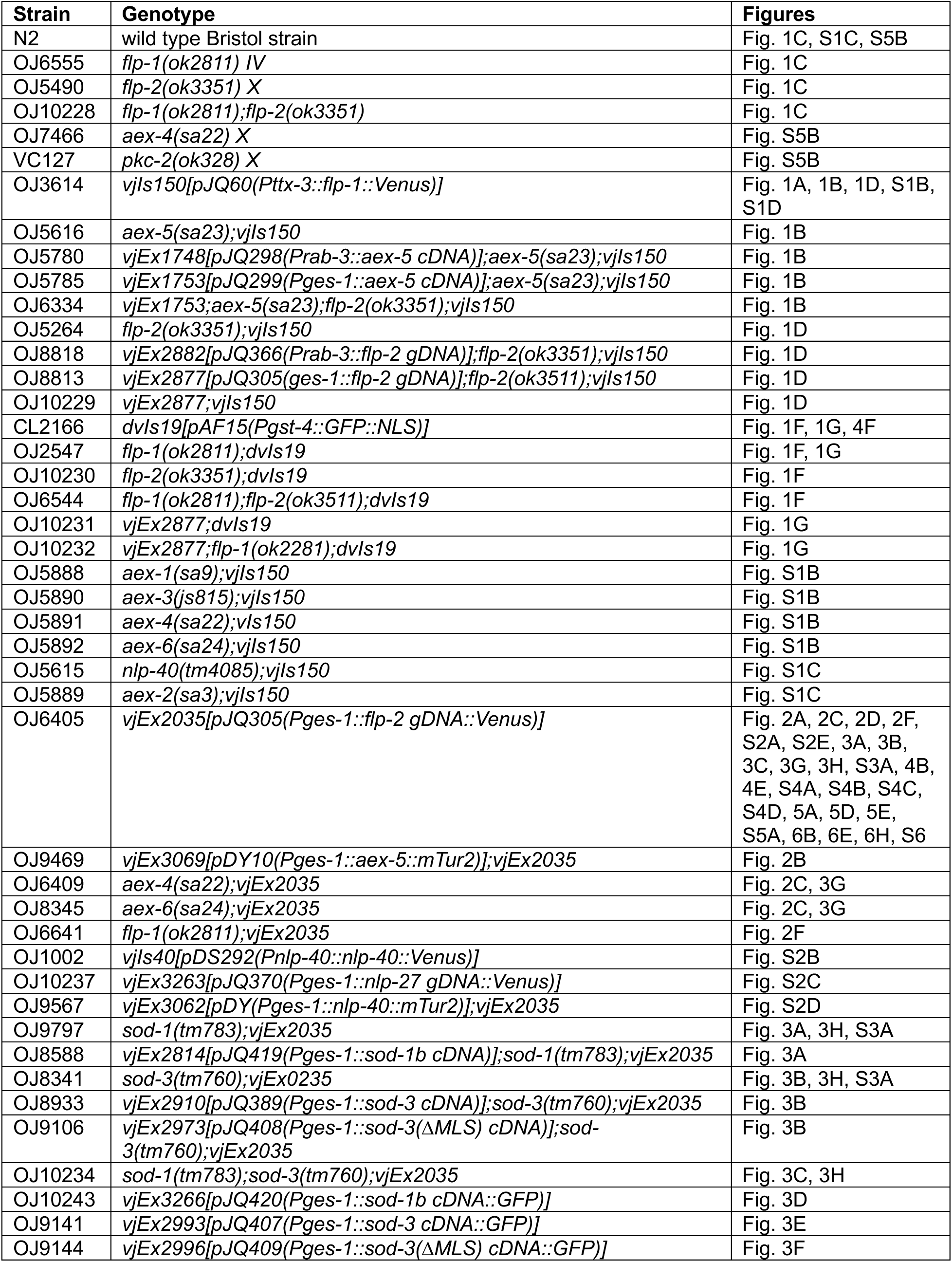
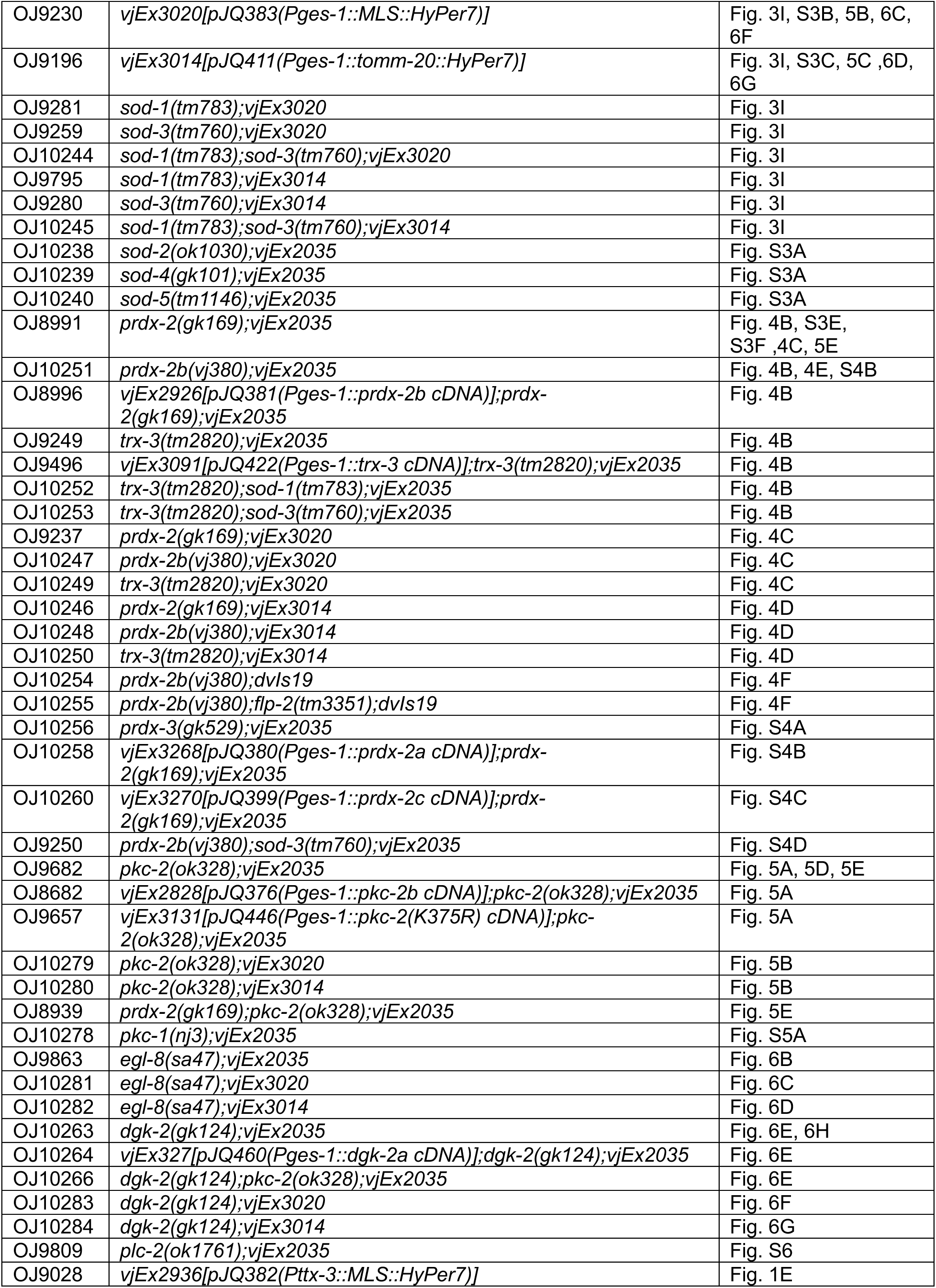
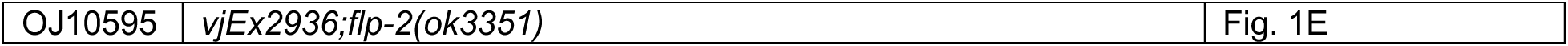
Strains, transgenic lines and plasmids used in this study.

